# Time-resolved tracking of cellulose biosynthesis and microfibril network assembly during cell wall regeneration in live Arabidopsis protoplasts

**DOI:** 10.1101/2024.07.29.605709

**Authors:** Hyun Huh, Dharanidaran Jayachandran, Junhong Sun, Mohammad Irfan, Eric Lam, Shishir P. S. Chundawat, Sang-Hyuk Lee

## Abstract

**Significance Statement:** Cellulose is a major extracellular matrix component of cells that is critical for plant development and has applications to bioenergy, agricultural food/feed, textile, and wood production. Cellulose is thought to be assembled by the closely coordinated motion of plasma membrane-embedded cellulose synthase enzyme complexes. To date, however, it has not been possible to visualize *de novo* plant cell wall synthesis at the single cell level with the necessary spatiotemporal resolution to derive a data-driven model of how plant cells can resynthesize and assemble cell wall after its removal. Based on our time-resolved data, we propose a new model for cellulose biosynthesis after successfully performing live protoplast time-lapse imaging to visualize for the first time the complex dynamics of *de novo* cellulose biosynthesis and assembly into an intertwined microfibril network.

Plant cell walls are composed of polysaccharides among which cellulose is the most abundant component. Cellulose is processively synthesized as bundles of linear β-1,4-glucan homopolymer chains via the coordinated action of multiple enzymes in cellulose synthase complexes (CSCs) embedded within the plasma cell membrane. Plant cell walls are composed of multiple layers of cellulose fibrils that form highly intertwined extracellular matrix networks. However, it is not yet clear as to how cellulose fibrils synthesized by multiple CSCs are assembled into the intricate cellulose network deposited on plant cell surfaces. Herein, we have established an *in vivo* time-resolved imaging platform for visualizing cellulose during its biosynthesis and assembly into a complex fibrillar network on the surface of *Arabidopsis thaliana* mesophyll protoplasts as the primary cell wall regenerates. We performed total internal reflection fluorescence microscopy (TIRFM) with fluorophore-conjugated tandem carbohydrate binding modules (tdCBMs) that were engineered to specifically bind to nascent cellulose fibrils. Together with a well-controlled environment, it was possible to monitor *in vivo* cellulose fibril synthesis dynamics in a time-resolved manner for nearly one day of continuous cell wall regeneration on protoplast cell surfaces. Our observations provide the basis for a novel model of cellulose fibril network development in protoplasts driven by complex interplay of multi-scale dynamics that include: rapid diffusion and coalescence of short nascently synthesized cellulose fibrils; processive elongation of single fibrils; and cellulose fibrillar network rearrangement during cell wall maturation. This platform is valuable for exploring mechanistic aspects of cell wall synthesis while visualizing cellulose microfibrils assembly.

## INTRODUCTION

Plant cell walls are complex extracellular matrices, which are predominantly composed of sink polysaccharides in the form of semi-crystalline cellulose microfibrils intertwined with amorphous matrix polysaccharides such as hemicellulose and pectin. They can be largely classified into primary and secondary cell walls that are assembled at different developmental stages with varying compositions and ultrastructural organizations of associated polysaccharides (1). The thinner and more extensible primary cell wall can better facilitate the continuous expansion of plant cells during growth and development. In contrast, the thicker secondary cell wall is more rigid and provides structural strength to plant cells and tissues (2). The primary cell wall, which is formed early in cell division and expansion, dictates the cell shape and acts as a template for the secondary cell wall deposition (3). The primary cell wall has been studied for a long time with particular emphasis on cellulose, which has been shown to form intricate fibrillar networks (4, 5). Cellulose, a linear β-1,4-glucan homopolymer chain, is synthesized by the plasma membrane-embedded glycosyltransferase enzyme family called cellulose synthase (CESA) that exists in various isoforms (6–8). CESAs self-assemble into a supramolecular protein complex called cellulose synthase complex (CSC) (8–12). Multiple cellulose chains are synthesized by a single CSC in a coordinated fashion involving extrusion out of the cell membrane to eventually form bundles of semi-crystalline cellulose fibrils, referred to as ‘elementary fibrils’, the smallest biological structural unit of cellulose (4, 13–15). Furthermore, multiple elementary fibrils can bundle into thicker fibrils, known as ‘cellulose microfibrils (CMF)’ (4, 15, 16).

Cell wall structure has been extensively studied by microscopy since the invention of the very first optical microscope by Robert Hooke, who described the boxlike cell walls of cork wood and other plants as ‘cells’ because they reminded him of cells in a monastery (17). More recently, electron microscopy (EM) has been used early on to visualize and study the structure of the fibrillar cell wall polysaccharides at higher resolution during plant cell development (18, 19). Atomic force microscopy (AFM) has also been widely used for studies of cell walls, because of its compatibility with imaging hydrated samples and minimal sample preparation requirements, contributing substantially to our understanding of plant cell wall structure (4, 15, 20–26). Both of these high-resolution microscopy techniques, while providing sharp nanoscale images, lack molecular specificity and are incompatible with live-cell imaging (4, 18, 19, 26). Fluorescence microscopy has significant advantages over other imaging methods in this regard, making it an important tool for visualizing distinct cell wall components. Plant cell wall-binding chemical dyes such as Calcofluor white (CFW), Congo Red, and Pontamine Fast Scarlet 4BS (PFS) are commonly used to stain cellulose in plant cell walls for fluorescence microscopy (27–35). However, most organic dyes bind nonspecifically to polysaccharides other than cellulose; may alter cellulose crystallinity *in vivo*; and can be toxic to living cells (36–38). In contrast, carbohydrate-binding modules (CBMs) are non-catalytic protein domains associated with the glycosyl hydrolases that bind to cell wall polysaccharides in a target-specific manner: CBM3a for crystalline cellulose (39–42); CBM17 and CBM28 for amorphous cellulose (42, 43); CBM76 for xyloglucan (44); and CBM27 for mannans (45). CBMs have been used to label various cell wall polysaccharides via conjugation with fluorescent proteins (23, 46, 47) or chemical dyes (35, 48–51). Such fluorescent probes can be imaged with widefield epifluorescence or scanning laser confocal microscopy (SLCM), but the diffraction-limited optical resolution often proves insufficient to resolve nanoscale fibril ultrastructures of cell wall polysaccharides. This diffraction barrier can be overcome with super-resolution fluorescence microscopy techniques (52–56), and such approaches have been applied to plants in recent decades (33, 57–67). However, application of super-resolution fluorescence imaging techniques to visualize cell wall polysaccharides is still in its early stages (33, 58, 59, 66, 67).

While we have a relatively good understanding of the mature plant cell wall structure, the dynamic processes by which polysaccharides are synthesized and assembled into fibrillar structures of distinct architectures within cell walls remain largely unknown. Real-time imaging of live plant tissues or cells with time-lapse microscopy is critical to bridge the knowledge gaps in our understanding of the mechanisms involved to construct a cell wall on the plant cell surface. In this respect, transgenic plants expressing CESA fusion to fluorescent protein (FP) have been utilized to visualize and track CESA/CSC motility in living plant tissues, and it has been shown that CESAs/CSCs move linearly at a speed of ∼0.2-0.4 μm/min, with their localization and motion guided by cortical microtubules and/or pre-existing cellulose microfibrils (68–77). CESA enzyme (or CSC) motion observed in such studies was presumed to be tightly coupled with cellulose synthesis, but direct high-resolution visualization of nascent cellulose fibril growth in plant tissues or isolated plant cells has not been reported yet, although cellulose fibril motion was captured in the context of reorientation during cell wall expansion (32), conformational change under tension (78), and degradation by hydrolytic enzymes (26). The polylamellate structure of plant cell wall is typically composed of ∼100 lamellae, with cellulose microfibrils forming a fine reticulated network on each lamellar surface (4, 18). Not surprisingly, it is very challenging to separately visualize only the growing nascent cellulose fibrils amid the background of underlying multilayered cellulose network on the surface of intact plant tissues. In that regard, plant protoplasts that could regenerate cell walls have tremendous potential to be utilized as a model system to study *de novo* cell wall synthesis *in vivo*. Confocal microscopy has been employed for 3D imaging of protoplasts (79–81), and even time-lapse imaging of cell wall regeneration was attempted for *Physcomitrella patens* protoplasts albeit at a low resolution (82). However, traditional confocal microscopy techniques have not proven suitable for long-term live-cell imaging of plant protoplasts owing to phototoxicity (83, 84). Furthermore, all previous protoplasts cell wall regeneration microscopy studies have used chemical dyes such as CFW and PFS to stain cellulose, which are not ideal for longer-duration, non-invasive, live-cell microscopy imaging that can lasts over several hours to days. Such limitations combined with the fragile nature of protoplasts, which makes them highly susceptible to damage by environmental/imaging conditions, have posed significant challenges for imaging the dynamic process of plant cell wall regeneration in real time.

In this work, we have succeeded to image cellulose biosynthesis and its assembly into CMF during the entire process of cell wall regeneration for Arabidopsis protoplasts and studied the processes through which intertwined cellulose fibril networks develop at the single cell level with high spatiotemporal resolution. To achieve this, we established an *in vivo* plant cell imaging platform based on Total Internal Reflection Fluorescence Microscopy (TIRFM), *in situ* cellulose fluorescence-labeling with fluorophore-conjugated tandem carbohydrate binding modules (tdCBM), and tight control of temperature and lighting conditions best suited for plant protoplast cell wall regeneration. We discovered that cellulose fibrillar networks on plant protoplast cell surfaces develop in multiple stages and our observations provide new insights into how cell walls are made in plants, which has important implications for developing transgenic crops that are optimized for cellulosic bioenergy and agricultural food/feed production.

## RESULTS

### Arabidopsis protoplasts regenerate cell walls in the presence of cellulose-labeling probes

Mesophyll protoplasts were isolated from leaves of 3–4 weeks old Arabidopsis Col-0 plants (Fig. 1a). We used medium-sized leaves, because they produced monodispersed protoplasts *versus* smaller or larger sized leaves. Upon incubation in a suitable plant protoplast cell wall regeneration media (WI/M2) under ambient white light provided by an LED bulb (Hue, Philips), protoplasts were biologically active and consistently regenerated cell walls for over 18 hours. They could be readily visualized by fluorescence microscopy using conventional calcofluor white (CFW) staining (Fig. S1, *SI Materials and Methods*). The experimental workflow is summarized in Fig. 1a.

**Figure 1:**
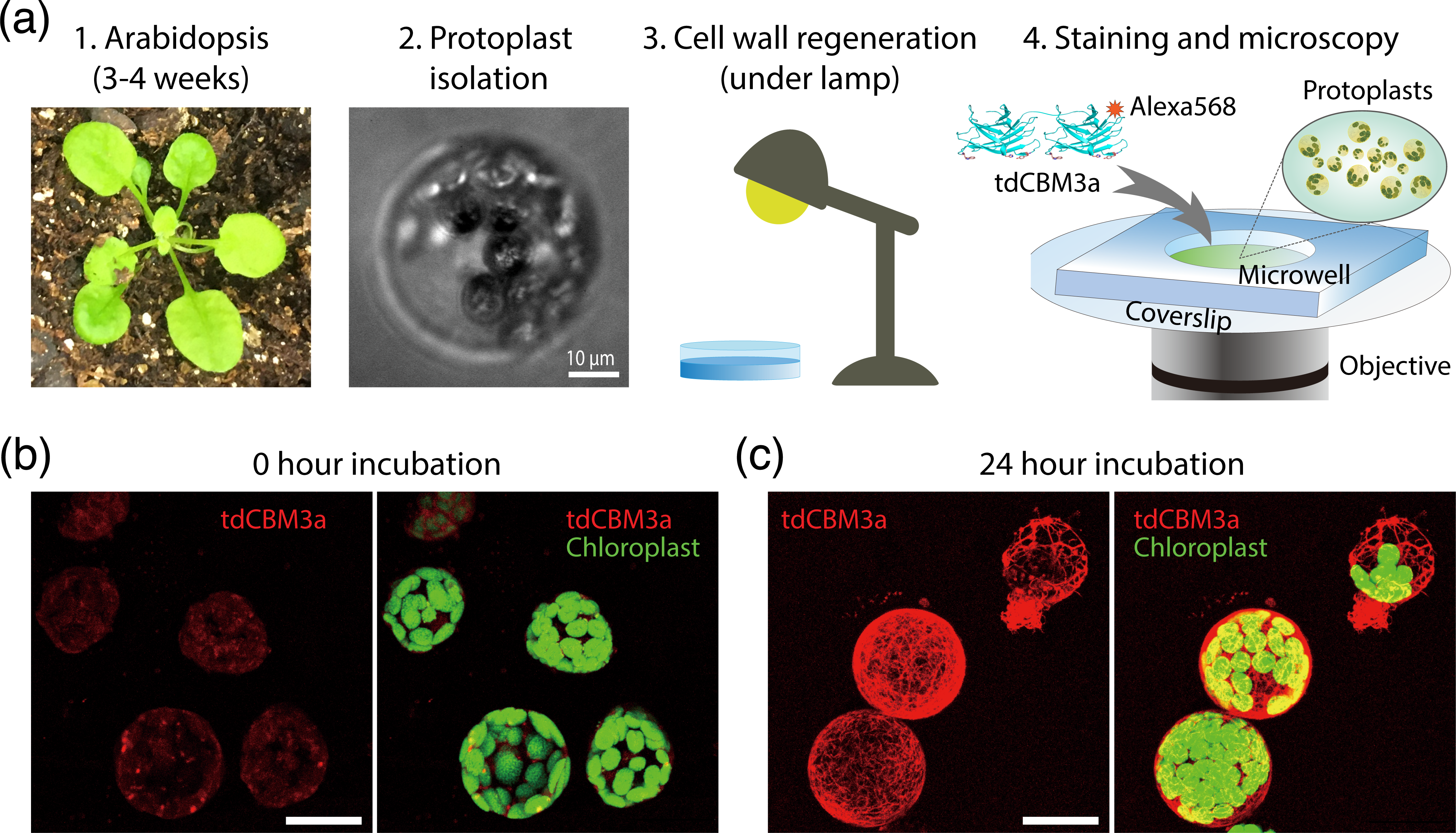
Workflow for imaging plant protoplast cell wall regeneration. (a) Summary of experimental workflow for imaging cellulose biosynthesis in protoplasts. Arabidopsis protoplasts were isolated from leaves of 3-4 weeks old plants by removing existing cell walls using enzymes and were incubated in a suitable cell wall growth media under a white light LED lamp. Fixed or live samples were stained with Alexa568-CBM3a to image regenerated cellulose fibrils formed on the protoplast cell surface. (b and c) Representative 3D confocal fluorescence micrographs of whole protoplasts with cellulose fibrils (red, pseudo-colored) and chloroplasts (green, pseudo-colored) visualized by Alexa568-CBM3a (561 nm excitation) and autofluorescence (488 nm excitation), respectively. Here, (b) shows images of cells prior to incubation with regeneration media and (c) shows cells after 24-hour incubation in cell wall regeneration media. Scale bars: 10 μm (a) and 20 μm (b-c).

CFW, although commonly used, is not optimal for visualizing cellulose in live cells because of its non-specific binding to various plant glycan epitopes and known cell toxicity (36–38). CBM-based proteinaceous probes were previously developed as a versatile, inert, and target-specific cell wall polysaccharide probe for enabling live-cell imaging as well as other immunolabeling applications (23, 35, 47–51, 39–46). In recent work, we developed a collection of engineered CBMs that have been extensively characterized by various biochemical and kinetic analyses to identify binding probes suitable for live-cell imaging. We determined tandem CBM3a (tdCBM3a) to be the most optimal CBM probe for imaging cellulose on Arabidopsis protoplast surfaces immersed in cell wall regeneration media (85) and further labeled it with Alexa Fluor 568 dye for long-term imaging (*SI Materials and Methods*). Using low concentrations of Alexa568-tdCBM3a (e.g., 100 nM) added to the cell wall regeneration media, we were able to continually label and image newly synthesized cellulose fibrils on the protoplast surface *in situ*, while maintaining high cell viability and low background fluorescence from unbound dyes over long incubation times. Protoplasts could develop dense cellulose microfibril networks after 24-hour cell wall regeneration under our experimental conditions (Fig. 1b-c), and showed no significant difference in the CMF architecture produced in the absence of the CBM probe (85).

### Cellulose is the major polysaccharide present in regenerated protoplast cell walls

While the Alexa568-tdCBM3a probe was anticipated to bind predominantly to cellulose (39–42), we further investigated whether the regenerated cell wall morphology visualized by the probe indeed represented cellulose networks by using cellulose enzymatic hydrolysis assays, glycome immunolabeling, and cell wall chemical composition analysis.

Protoplasts with regenerated cell walls were fixed with glutaraldehyde to preserve cell wall integrity and arrest cell wall development after 24 hours. Cells were then labeled with Alexa568-tdCBM3a for visualizing the regenerated CMF network. When fixed protoplasts were subject to hydrolysis with cellulase cocktail (Cellic CTec2, Novozymes) (86), we observed complete disappearance of fibrillar structures of the regenerated cell walls within 40 minutes (Fig. 2a, *SI Video 1*). Although this result documented that cellulose was present within the fibrillar structures, comprising the visualized fibril network pattern, we could not exclude the possibility that we were detecting loss of xyloglucan labeled with tdCBM3a (87) instead of cellulose. (*Note: CTec2 cellulase cocktail is known to contain endoglucanases capable of hydrolyzing xyloglucan and cellulose*). To examine this possibility, we immunolabeled xyloglucan with xyloglucan-specific antibody (LM15) that was conjugated with anti-rat IgG Alexa Fluor 488 secondary antibody (88, 89). Two-color confocal microscopy with Alexa568-tdCBM3a and Alexa488-LM15 indeed showed almost identical regenerated protoplast cell wall morphology in the two fluorescence channels (Fig. 2b). However, when a purified cellulose-specific hydrolytic enzyme (E-CELBA, Megazyme) with no detectable activity towards xyloglucan was added, Alexa568-tdCBM3a fluorescence completely vanished whereas Alexa488-LM15 fluorescence still remained. Interestingly, the Alexa488-LM15 fluorescence signal in the presence of E-CELBA lost the fibril network pattern and instead was more homogeneously distributed over the cell surface (Fig. 2b). Taken together, these results strongly suggest the cellulose fibril-specific binding capacity of tdCBM3a and the potential for close association between cellulose fibrils and xyloglucan — the major hemicellulose of plant primary cell walls — to be present on regenerated protoplast cell wall surfaces (90).

**Figure 2:**
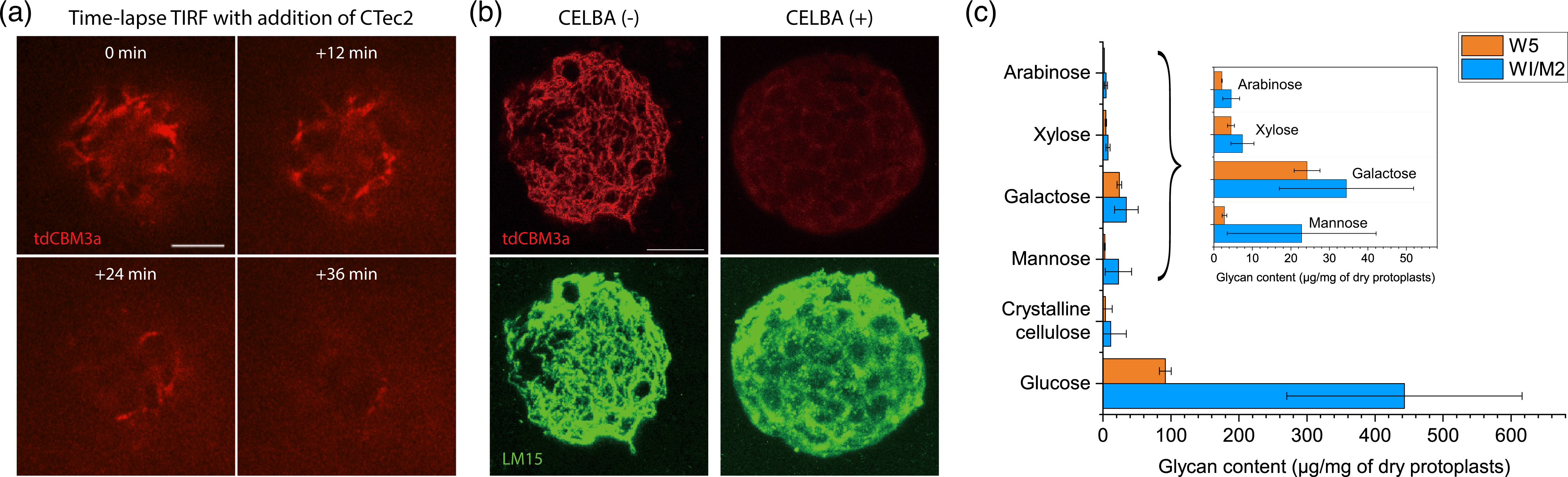
Identification of cellulose on regenerating protoplast cell membrane surfaces. (a) Time-lapse TIRFM images of a protoplast cell surface with its regenerated cell wall fibrillar network being continuously degraded by a commercial cellulase (Cellic CTec2, Novozymes). The fibrillar network on the cell was stained with Alexa568-tdCBM3a. The initial time point (0 min) represents the time at which the CTec2 cellulase was added. (b) Two-color confocal images of regenerated protoplasts cell walls and fibrillar network that were labeled with Alexa568-CBM3a (red, pseudo-colored) and Alexa488-LM15 (green, pseudo-colored) to visualize cellulose and xyloglucan, respectively. Left panel: Cell surface showed patterns of fibril network that were colocalized between the two labels. Right panel: Addition of cellulose-specific hydrolytic enzyme (E-CELBA) had Alexa568 fluorescence signal completely disappear for all observed protoplasts; whereas the Alexa488 fluorescence remained strong even after the fibril network pattern disappeared. (c) Gas chromatography-mass spectrometry (GC-MS) analysis of monosaccharides released after acid hydrolysis of protoplast samples to analyze the chemical composition of cell wall polysaccharides that were produced after incubating protoplasts for 24 hours in the regeneration media (WI/M2) in comparison to the control media (W5). Inset: Magnified view of arabinose, xylose, mannose, and galactose data. Data are represented as mean and standard deviation values that are obtained from n=3 technical replicates. Scale bar: 5 μm (a) and 10 μm (b).

We next performed acid hydrolysis followed by gas chromatography-mass spectrometry (GC-MS) on plant protoplasts with regenerated cell walls to determine the precise chemical composition of matrix polysaccharides (Fig. 2c). Protoplasts with cell walls regenerated for 24 hours in a suitable regeneration media (WI/M2) showed ∼4.5-fold increase in the amount of glucan polymers as compared to protoplasts lacking cell walls (incubated in W5 media as control). Sugar composition estimated via GC-MS analysis was performed subsequent to acid hydrolysis of the polysaccharides into their respective monosaccharides, and therefore the glucan content reported here could have been partially derived from both amorphous cellulose (*β*-1,4-glucan) and callose (*β*-1,3-glucan). During the initial stages of cell wall regeneration, callose is known to interact with cellulose to provide stiffness and support (91, 92). However, callose is mainly found in pollen tubes, plasmodesmata, and wounded cells, whereas cellulose is the most abundant structural polysaccharide constituting 30-90% of plant cell walls (93). Therefore, in light of the cellulase activity and tdCBM binding specificity results, the large amount of glucan estimated during GC-MS is mostly indicative of presence of cellulose as the major polysaccharide component in regenerated cell walls. In addition, other minor polysaccharides composed of other types of monomer sugars such as galactose, mannose, or xylose were also detected and showed slightly increased amount in the protoplasts with regenerated cell walls although in much lower absolute amounts than glucan polymers (Fig. 2c, Inset).

Interestingly, crystalline cellulose (as determined to be a fraction of glucan polymers resistant to mild but not to concentrated sulfuric acid hydrolysis prior to GC-MS analysis) was also detected but only in small quantities (Fig. 2c). It is expected that amorphous or semi-crystalline cellulose would be susceptible to even mild acid hydrolysis to produce glucose. In contrast, crystalline cellulose, while being recalcitrant to mild acid treatment, would be hydrolyzed by exposure to concentrated sulfuric acid. Therefore, our cell wall composition results suggest that the nascent cellulose fibrils, as visualized by fluorescence microscopy (Fig. 1c, Fig. 2a-b), likely contain a significant fraction of amorphous or semi-crystalline cellulose. To gain further validation into the crystallinity of the nascent cellulose on regenerated protoplast cell walls, we labeled CBM17 (a carbohydrate binding module specific to amorphous cellulose) with Alexa Fluor 568, making Alexa568-CBM17. In contrast to the original CBM3a probe developed to bind to both crystalline and semi-crystalline cellulose (39–42, 94), CBM17 is known to bind exclusively to amorphous or semi-crystalline cellulose (42, 43). The protoplasts stained with Alexa568-CBM17 after ∼25-hour cell wall regeneration showed fibril network structure that looked very similar to the protoplasts stained with Alexa568-tdCBM3a (Fig. S2). We also tested CBM1, known to bind to crystalline cellulose exclusively (95–97), and found that Alexa568-CBM1 did not show any fluorescence (Fig. S2). These results imply that mostly amorphous or low crystallinity cellulose is likely to be predominantly present in the nascently synthesized and assembled cellulose microfibrils on plant protoplast surfaces after cell wall regeneration under our experimental conditions.

### Live-cell time-lapse protoplast imaging enables near real-time visualization of nascent cellulose biosynthesis

Live-cell time-lapse fluorescence microscopy is essential for us to gain full understanding of the dynamic process of how the mature cellulose fibril network (as seen in Fig.1c) develops over longer time periods (∼24 hours) of cell wall biosynthesis at the single cell level. Despite the extensive microscopy studies conducted on plant cells and tissues over the many decades (4, 15, 18–26, 32–35, 46–51, 57–81), continuous visualization of cellulose on the surface of isolated plant protoplast cells during cell wall regeneration has been rarely reported (82). This paucity of data can be primarily ascribed to the challenge in keeping plant protoplasts healthy or viable over a prolonged imaging period, because protoplasts lacking cell walls are often susceptible to mechanical, chemical, and light-induced damages. We employed two main strategies for reducing protoplast stress during longer-term live-cell fluorescence microscopy, including 1) *in situ* cellulose labeling with a low concentration of CBM-based probes like Alexa568-tdCBM3a; and 2) imaging the underside of the protoplast with Total Internal Reflection Fluorescence Microscopy (TIRFM) for minimal dose of excitation light and phototoxicity. The TIRFM excitation scheme was optimized to reduce background fluorescence from unbound dyes in solution and autofluorescence from chloroplasts.

We additionally found that environmental conditions, especially ambient light and temperature, substantially influenced cell wall regeneration, and thus, proper control of these environmental conditions over the entire course of imaging was critical. Protoplasts incubated without ambient light failed to regenerate cell walls, demonstrating the essential requirement for light for cell wall regeneration (Fig. S3). To provide protoplasts with controlled ambient light during time-lapse fluorescence microscopy, we synchronized a programmable white LED light bulb (Hue, Phillips) with our microscope such that the bulb automatically switched off and on at the start and end of an image acquisition cycle, respectively. Initially, we found that cell walls were not properly regenerated when protoplasts were incubated on our microscope stage even in the presence of ambient light. We found that the motorized microscope body and stage generated heat (increasing sample temperature to 25 °C or higher). We, therefore, developed an active liquid-cooling system for the microscope stage and objective lens to maintain the sample chamber temperature at ∼18 °C) (Fig. 3a). These modifications eliminated inconsistent cell wall regeneration under longer-term imaging conditions.

**Figure 3:**
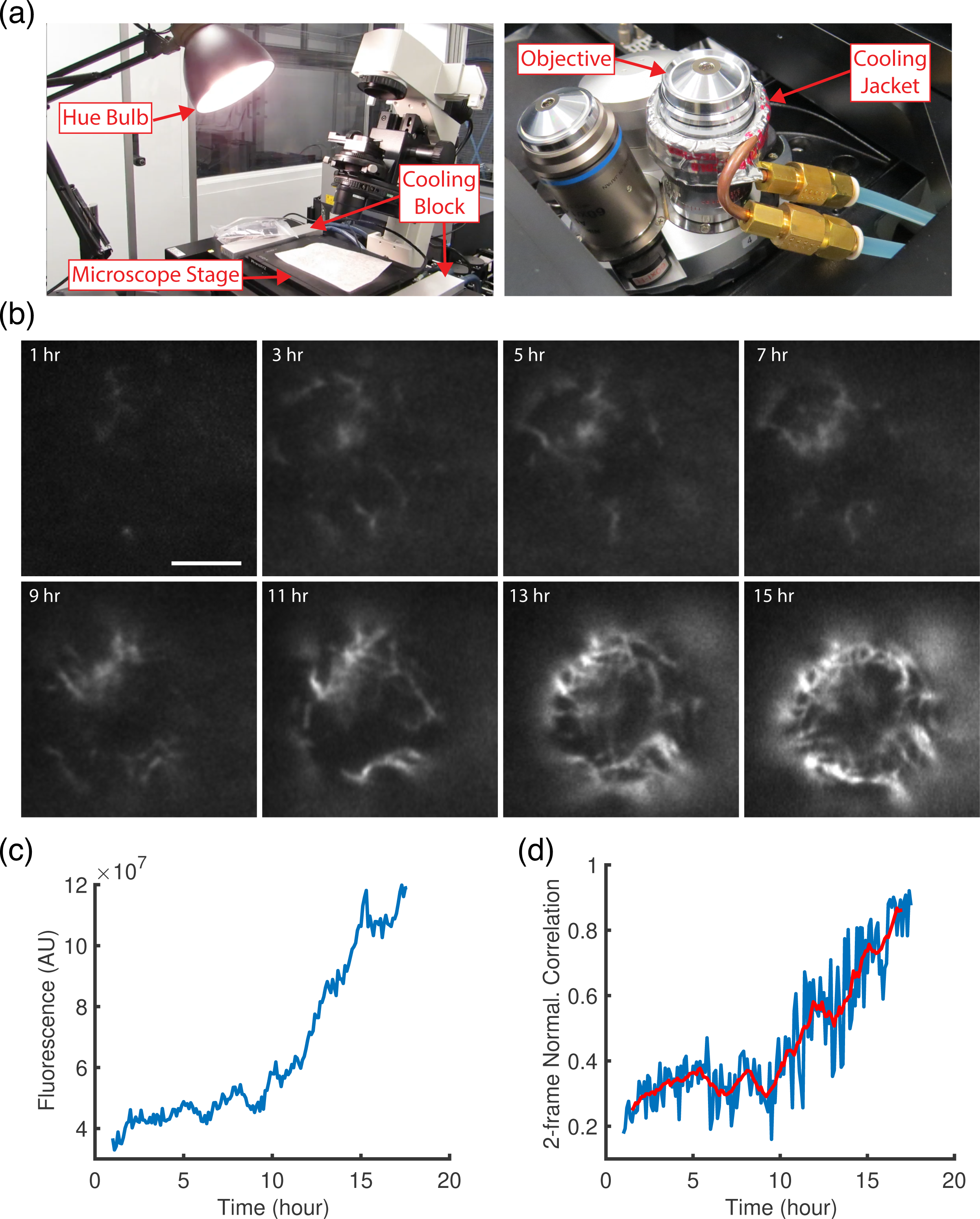
Live-cell TIRFM to visualize real-time cellulose biosynthesis in protoplasts. (a) Pictures of the custom-built imaging instrument parts to control the environmental conditions, i.e., ambient light (left) and sample temperature (right), while acquiring real-time TIRFM images of cellulose that are being synthesized and assembled on protoplast cell surfaces. (b) Representative time-lapse TIRFM image sequences of cellulose formation on a protoplast surface. Cellulose was *in situ* stained with Alexa568-tdCBM3a in the cell wall regeneration media. Image acquisition began 1 hour after the start of cell wall regeneration and continued for ∼16 hours at 6-minute imaging interval. Images subsampled at 2-hour interval are shown here (see *SI Video 2* for the full image stack). (c and d) Total fluorescence (c) and normalized cross-correlation (d) in time for the time-lapse image data corresponding to *SI Video 2*. Total fluorescence and normalized cross-correlation (NCC) (see *SI Materials and Methods* for definition) quantitatively estimate the total amount and the overall mobility, respectively, of cellulose on the protoplast surface that can be observed by TIRFM. A lower NCC value implies a higher fibril mobility and vice versa. Blue curves (in c and d) represent the original 6-miniture interval time-lapse data. Red curve (in d) represents the moving average of the blue curve with a sliding window of 1-hour width. Scale bar: 5 μm.

TIRFM, combined with CBM-based *in situ* labeling and active control of ambient light and temperature, enabled us to visualize and image cellulose growth on the underside of the protoplast every 6 minutes continuously for over 16 hours (Fig. 3b, *SI Video 2*). Interestingly, the time-lapse images showed that the amount of nascent cellulose increased nonlinearly in time, with an initial ‘lag’ phase—lasting hours (0-10 hours)—followed by a shorter ‘rapid growth’ phase (Fig. 3c). Highly mobile, short (≲ 2 μm) cellulose fibril fragments were mainly present in the lag phase, whereas less mobile or stationary networks of longer intertwined cellulose fibrils were predominantly observed during the rapid growth phase. To quantify overall mobility of cellulose fibrils *versus* time, we plotted normalized cross-correlation (NCC) between two consecutive image frames over time (Fig. 3d, *SI Materials and Methods*). NCC analysis confirmed an interconnection between the mobility and the amount of nascent cellulose fibrils in a biphasic manner (Fig. 3c-d). High fibril mobility (*i.e*., low NCC) visible during the early lag phase rapidly decreased (*i.e*., increased to high NCC) in the later rapid growth phase of cellulose production.

### Nascently synthesized short cellulose fibrils move diffusively on the protoplast surface

The image acquisition with 6-minute time interval used for longer-term TIRFM proved too slow to reveal the details of cellulose fibril dynamics on the surface of the protoplast during distinct stages of cell wall regeneration. When the image acquisition time interval was shortened to once every 20 seconds, the increased total dose of excitation laser exacerbated phototoxicity of cells and reduced cell viability, preventing longer-term live-cell imaging. As a compromise, we imaged different cell wall regeneration stages for only several hours at 20-second intervals to better understand the dynamics of cellulose network development during different phases of regeneration without overstressing the protoplasts.

20-second interval time-lapse imaging showed random translational and rotational motion of very short cellulose fragments on the protoplast underside during the early stages of cell wall biosynthesis (Fig. 4a, *SI Video 3*). We selected short (≲ 2 μm) fibril fragments, manually tracked their centers in two dimensions, and plotted their mean squared displacement (MSD) in time. Fitting the MSD of each trajectory with a power-law scaling in time resulted in a unimodal distribution of the scaling factor ‘α’—also known as anomalous diffusion exponent—with 0.89 (± 0.05, SE) as the fitted mean value (Fig. 4b). Therefore, the cellulose fragments formed at the early stages of cell wall biosynthesis moved rapidly on the protoplast cell surface according to nearly normal diffusion (i.e., α ≃ 1), if not slightly sub-diffusive. The closed topology of the spherical cell surface may be at least partially responsible for sub-diffusive 2D motion, when the cellulose fibril motion on the 3D spherical surface is projected onto a 2D focal plane in TIRFM imaging.

**Figure 4:**
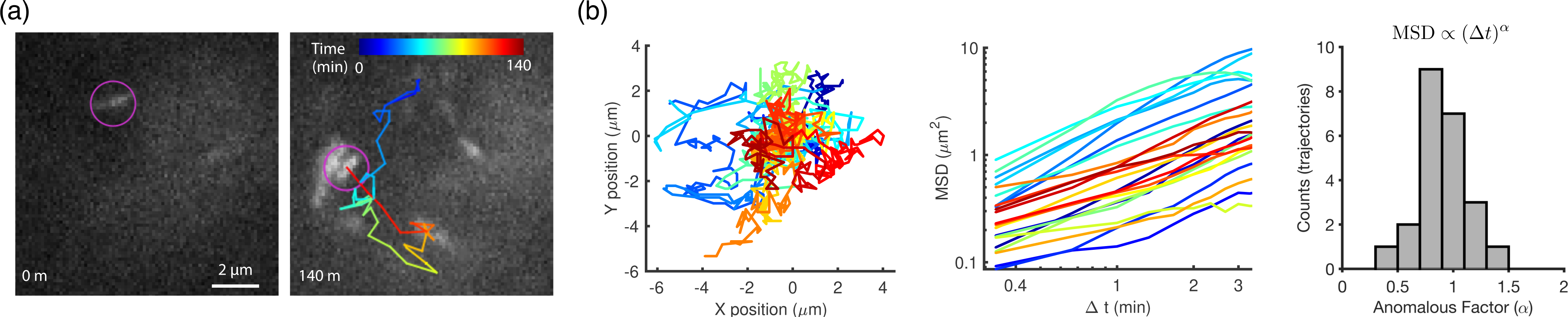
Diffusive motion of short cellulose fragments can be seen in the early cellulose biosynthesis and cell wall regeneration stage. (a) Trajectory of the center for a representative short cellulose fragment moving rapidly on the surface of an Arabidopsis protoplast at the early stage of cell wall regeneration. The colormap represents the tracking time (see *SI Video 3* for the full image stack). Cellulose was imaged on the surface of regenerating Arabidopsis protoplasts through *in situ* staining with Alexa568-tdCBM3a and TIRFM. (b) Left: 2D trajectories of 23 short fibrils (≲ 2 μm in length) with respective motions that looked like random walks. The trajectories were colored arbitrarily and centered at the origin of XY-axes for better visualization. Middle: Log-log plot of mean squared displacement (MSD) in lag time for the 23 trajectories. The temporal scaling factor of MSD (i.e., anomalous diffusion exponent) was obtained by linear fitting of this plot. Right: Histogram of anomalous diffusion exponent for the 23 trajectories (mean=0.89, standard deviation=0.24). Scale bar: 2 μm.

### Short/thin cellulose fragments can coalesce into longer/thicker fibrils to form a cellulose fibril proto-network

The short cellulose fibril fragments were not only diffusing on the membrane surface but could also change shape through rapid inter-fibrillar interactions. Figure 5a (also see *SI Video 4*) exemplifies such cellulose inter-fibril coalescence events. It was frequently observed that two short fibrils (labeled ‘1’ and ‘2’ in Fig. 5a) of ∼2 μm or less in length encountered each other through surface diffusion. Typically, an end of one fibril first made a contact with the middle part of the other fibril obliquely and they soon collapsed on top of each other, creating a thicker fibril (labeled ‘3’ in Fig. 5a). Interestingly, the bundle of two coalesced fibrils changed its morphology. For example, the short, thick bundle (‘3’ in Fig. 5a) later emerged as a longer, thinner fibril possibly through sliding motion between the two merged fibrils (i.e., ‘1’ and ‘2’). Sliding of cellulose fibrils was observed in previously published AFM studies and proposed to play an important role in building extensible cell walls (5, 78, 98). Inter-fibril coalescence also occurred among longer cellulose fibrils (see ‘3’, ‘4’, and ‘5’ labels in Fig. 5a) to generate even longer fibrils. In this case, one fibril could even swing around, pivoting on the junction with another fibril until their association was stabilized. Based on such observations, we reasoned that the cascade of cellulose fibril coalescences occurring at various fibril length scales would contribute toward building an intertwined cell wall fibrillar framework for less mobile, extended fibril networks. In addition, coalesced cellulose fibrils sometimes showed highly dynamic reshaping of their morphology (Fig. 5b, *SI Video 5*). Cellulose fibrils can also exhibit remarkable dynamics of bending and unbending (Fig. 5c, *SI Video 6*) such that a fibril even self-coalesced through ∼180° bending (*SI Video 7*). It was apparent that the dynamic morphological plasticity and the ‘sticky’ non-covalent interactions between cellulose fibrils could play important roles in driving the self-assembly of a sparsely interconnected primordial cellulose fibril network observed herein, which we termed “proto-network”, as a potential precursor to the final mature, dense cellulose fibril network.

**Figure 5:**
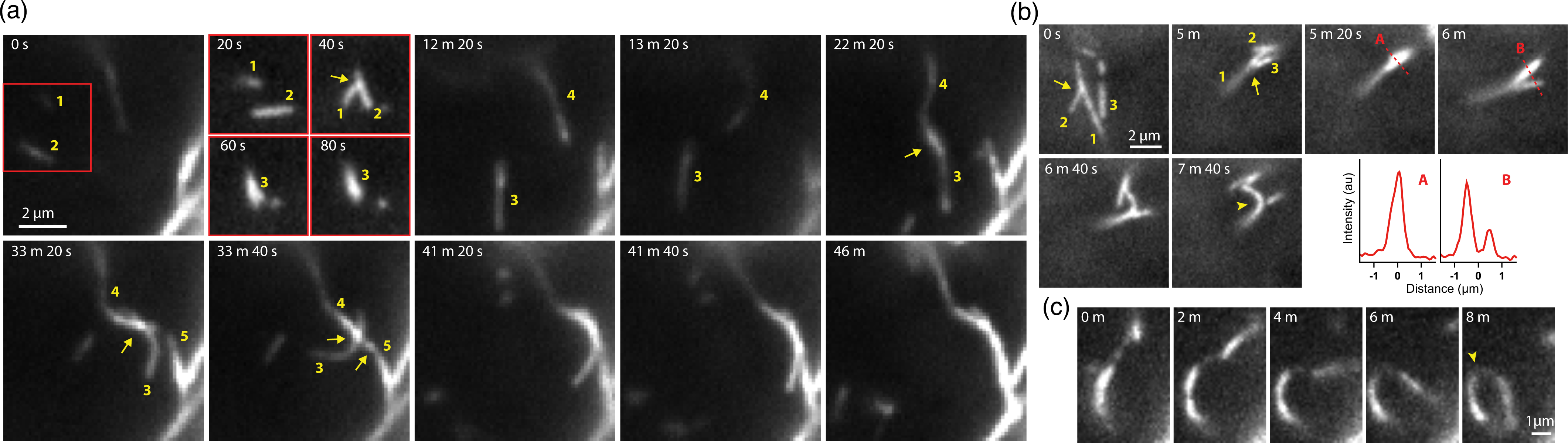
Coalescence of nascently synthesized cellulose fibrils can lead to generation of thicker and/or longer fibrils on the protoplast surface. (a) Time-lapse images showing a cascade of inter-fibril coalescence events that led to assembly of a stable, extended, and thicker cellulose fibril. These images are subsampled from *SI Video 4* that was acquired at 20-second interval. The second panel shows four image sequences of the subarea in the first panel indicated by a red square. Relevant fibrils are numbered in yellow to identify coalescence events, e.g., ‘1’ and ‘2’ fibrils coalesce into ‘3’. (b) An example of coalesced cellulose fibrils showing dynamic morphological changes (see *SI Video 5* for the full image stack). (c) Images of an extended cellulose fibril showing dynamic bending and unbending (see *SI Video 6* for the full image stack). The yellow arrow (in b) indicates the junction of fibril coalescence. The yellow arrowhead (in b and c) indicates the point of high curvature. Cellulose was imaged on the surface of regenerating Arabidopsis protoplasts by *in situ* staining with Alexa568-tdCBM3a and TIRFM.

### Processive elongation of cellulose fibrils further crosslinks and densifies the cellulose network on the protoplast surface

Cellulose fibrils are thought to be synthesized by mobile CSCs on cell membrane surfaces (9–12), and as such we expect to observe actively growing fibrils on the protoplast surface. However, we rarely observed such events in the early stages of cell wall biosynthesis (< 10 hours) when only highly mobile short fibril fragments were prevalent. As described previously, small fibrils were rapidly diffusing, tumbling, and frequently disappearing out of focus, which made it technically difficult to track short fibril length changes. It was not until the emergence of a proto-network (∼10 hours after the start of cell wall regeneration) that distinct processive synthesis, elongation, and deposition of single fibrils were clearly observed. *SI Video* 8 and Figure 6a show such events of single fibril biosynthesis observed in a protoplast over ∼7-hours after 16-hour incubation for cell wall regeneration. Such newly synthesized fibrils were often seen to be anchored to the relatively immobile proto-network on one end while their length grew processively on the other end. Diffusive motion was still apparent in the actively growing fibrils, but the active adherence of the growing fibrils to a stable underlying proto-network substantially reduced the random motion as compared to freely diffusing short cellulose fragments seen otherwise (Fig. 6b). A single growing fibril sometimes encountered another growing fibril, and they both fused to form a bridge between two anchor points on the existing fibril network (Fig. 6c). Fibrils also grew and crossed over other stationary, formerly deposited fibrils, creating a point-like coalescence at each junction (Fig. 6d). Therefore, active growth and processive deposition of single fibrils played an important role in interconnecting and crosslinking the underlying proto-network to develop an increasingly dense and stable cellulose fibril network on the protoplast surface. Based on our time-lapse microscopy data, the processive growth rate of a single fibril was estimated to be ∼0.13 (± 0.04, STD) μm/min (Fig. S4). This rate, although slightly slower, is in the similar range as the previously reported speed of CESA/CSC (∼0.20-0.40 μm/min) movement measured in plant tissues that was supposedly coupled to cellulose synthesis (68–76).

**Figure 6:**
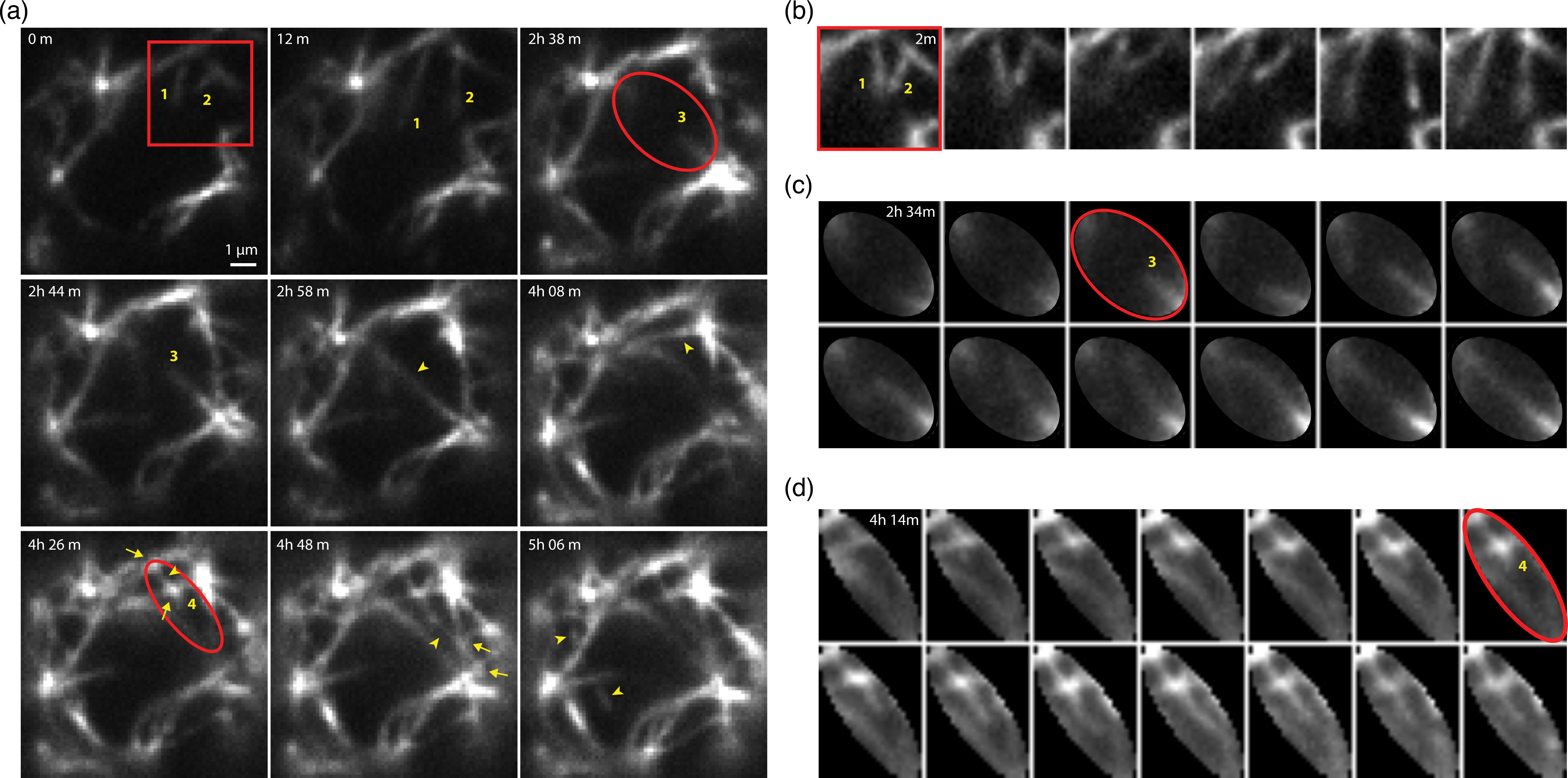
Observation of real-time processive biosynthesis of single cellulose fibrils on the protoplast surfaces using TIRFM. (a) Time-lapse images showing processive growth of single cellulose fibrils on the protoplast surface (see *SI Video 8* for the full set of images). The image acquisition started after 16-hour cell wall regeneration. The representative growing fibrils are marked by yellow numbers. Yellow arrowheads indicate the fibril segments that newly appeared upon processive fibril synthesis and extension. Yellow arrows indicate the coalescence junctions newly formed by actively growing fibrils that cross over underlying preexisting fibrils. (b-d) Detailed image sequences of three subareas in Fig. 6a that are highlighted by a red box and two ovals. The time interval between two consecutive images is 2 minutes in b, c, and d.

### Cellulose proto-network evolves its morphology toward a more compact and stable state at later stages of cell wall regeneration

We observed that the ultrastructure of the newly assembled proto-network continued to change even after its emergence (Fig. 7, *SI Video 9*). Certain local areas of the proto-network, especially where the overall matrix mesh was relatively coarse, appeared more mobile and unstable than other areas (Fig. 7b). Fibrils in these areas showed pronounced diffusive motion and bending dynamics; and furthermore, they could deform to a more compact appearance by mechanical pressure from the neighboring parts of the fibril network that could be more rigid (Fig. 7a). Normalized cross-correlation or NCC analysis plots of the time-lapse images implied that such densification and refinement of the fibril mesh ultimately results in a more stationary, stable cellulose networks (Fig. 7c). Therefore, the cellulose fibrillar network appears to be evolving slowly (*i.e*., over more than 10 hours) but continuously to form a more compact and stable multi-layered structure on the protoplast membrane surface.

**Figure 7:**
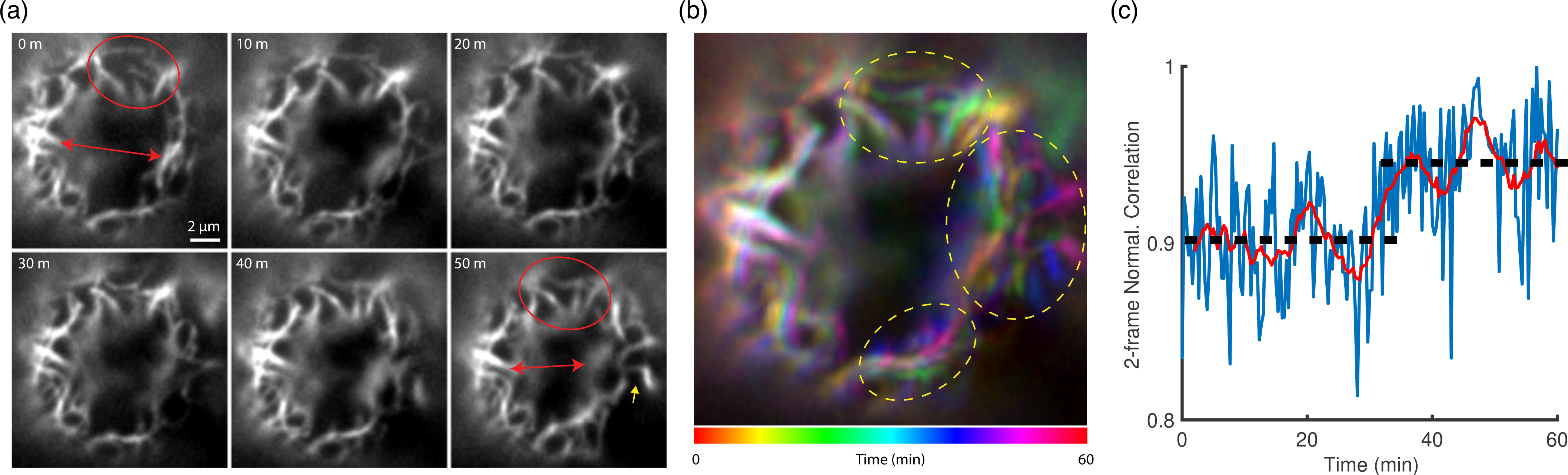
Cellulose fibril network continues to mature and reshape on the cell surface. (a) Time-lapse images of a cellulose fibril network with a morphology that slowly but continuously changes as the cell wall regeneration process goes to completion (see *SI Video 9* for the full set of images). The initial coarse mesh of fibrils within a highlighted area (red oval) turned into a more refined mesh within 50 minutes. The initial distance between two specific locations on the fibril network (red arrow) got shorter by intrusion of another fibril segment indicated by yellow arrow (see *SI Video 9* for the detail). (b) 2D projection of the time-lapse images with temporal color coding to visualize the local mobility of fibril network. Mobile regions appear in rainbow color (dotted ovals) whereas stationary regions in white color. (c) Normalized cross-correlation shows a trend of slow but increasing values in time, implying evolution of the network towards a more stable ultrastructural configuration. Blue curve represents the original 20-second interval time-lapse data. Red curve represents the moving average of the blue curve with a sliding window of 4-minute width. Scale bar: 2 μm.

### Super-resolution microscopy reveals intricate cellulose fibril network structures deposited in the newly regenerated protoplast cell wall

The distinctive six-lobed rosette structure of CSC, with each lobe composed of a CESA homotrimer, implies that 18 cellulose chains are likely to assemble and bundle into a cellulose elementary fibril (9–12). AFM and other imaging methods have indeed measured ∼3-5 nm diameters of elementary fibrils, which is consistent with cellulose microcrystal fibril packing models of 18, 24, or 36 cellulose chains (4, 13–15). Moreover, elementary fibrils are thought to merge into bundles with various diameters of less than 100 nm, termed ‘microfibrils’ (4, 15, 16). Because of the diffraction-limited optical resolution, it was unclear whether our TIRFM images of cellulose fibrils on the protoplast surface represented elementary fibrils, or microfibrils, or even bundles of microfibrils. To visualize the cellulose fibril network in higher resolution, we applied stochastic optical reconstruction microscopy (STORM) (55), a super-resolution fluorescence microscopy technique in the category of single-molecule localization microscopy (SMLM) (54–56). We found that Alexa568-tdCBM3a “blinked” very well under STORM imaging conditions, which is the most critical requirement for identifying and localizing single molecules in SMLM (*SI Video 10, SI Materials and Methods*). While we were trying to immobilize live protoplast cells— large spherical objects of 40-50 μm in diameter—on a poly-L-lysine-coated coverslip for STORM imaging, the entire cellulose fibril network was found to be susceptible to detachment from the cell surface through hydrodynamic drag forces. We serendipitously discovered that during detachment, protoplasts peeled in a way as to leave behind a wide area of the cellulose fibril network on the poly-L-lysine coated coverslip surface. We further developed a procedure to optimally imprint a large area of cellular surface cellulose network on the coverslip, while largely preserving native structure (Fig. S5A). The cellulose-visible area of ∼35 μm diameter implied that the cellulose fibrils network covering approximately a quarter of the original protoplast surface was readily transferred to the coverslip. This was a much larger area than the ∼10 μm diameter area of the protoplast bottom surface that could be typically imaged by TIRFM in focus. The imprinted cellulose fibril network looked similar to those of intact cells and we postulated that 2D-STORM images of the imprinted cellulose could provide insights into the high-resolution architecture of regenerating protoplast cellulose fibril network while avoiding the challenges in 3D-STORM imaging of thick objects such as intact plant protoplast cells (99, 100).

High-resolution STORM images revealed an intricate reticulated network of cellulose fibrils on the protoplast surface after 48-hour cell wall regeneration in the presence of our Alexa568-tdCBM3a label (Fig. 8). Single fibrils in STORM images appeared as much thinner lines (≲ 50 nm) than in TIRFM images (Fig. 8a). A seemingly single fibril in TIRFM images was frequently found to actually consist of multiple fibrils in STORM images (Fig. 8b, Fig. S5b). A layer of cellulose fibrils in plant cell walls are often seen to orient in a certain direction, as in the case of onion epidermis, Arabidopsis petioles/hypocotyl/roots cells, and other plant systems (4, 23, 26, 32, 33). In contrast, the fibrils seen in our STORM images did not show any seemingly preferred orientation in regenerating protoplasts, but instead formed a highly reticulated network like knitted cloths, fish nets, or trellis. Fibril density or mesh size in the network was heterogeneous, with the fibrils in coarse mesh areas appearing as more curved than in fine mesh areas (Fig. S5b). We further identified various types of inter-fibril associations in the assembly of the fibril network (Fig. 8c-d, Fig. S5b). A junction between two crossing fibrils was the prevalent type (arrow symbol in Fig. 8c-d and Fig. S5b), which was likely to be formed by an actively growing fibril deposited across an underlying stationary fibril as seen in the case of Fig. 6d. It was also common for the convex side of a curved fibril to tangentially coalesce with another straight or curved fibril (two-headed arrow symbol in Fig. 8c-d and Fig. S5b). In this case, the fused segment appeared as a single, brighter line within the STORM resolution limit. Such an organization might have resulted from a lateral encounter and fusion between two dynamically bending/unbending long fibrils that were either freely diffusing (Fig. 5) or tethered to the underlying network (Fig. 7). Interestingly, STORM images could differentiate two very close but non-coalesced fibrils (chevron symbol in Fig. 8c-d and Fig. S5b) from the truly fused or coalesced ones. The motion of such closely juxtaposed but non-coalesced fibrils might have appeared as reversible coalescence events seen in the lower resolution TIRFM images or movies (Fig. 5b). In some cases, two fibrils remained closely apart (< 200 nm) over ∼1.5 μm segment between the fused ends on both sides, resembling a highly stretched loop (Fig. 8e). We also observed similar cellulose fibril bifurcation patterns under scanning electron microscopy (Fig. 8f). Typically, two fibrils branched out of a bundled segment, but branching of even more fibrils was also observed (see dotted circle in Fig. S5b).

**Figure 8:**
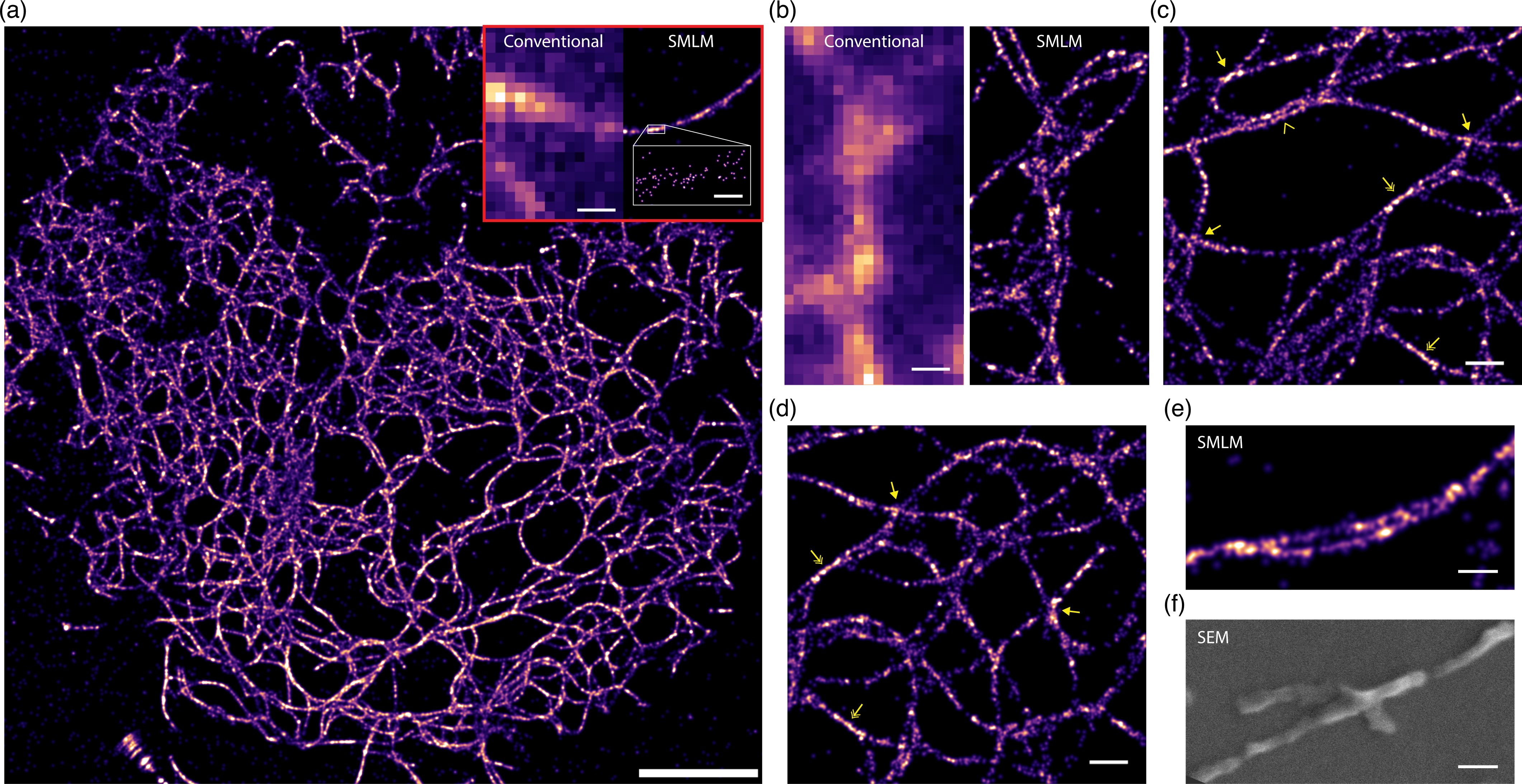
Super-resolution 2D-images of the protoplast cellulose fibril network imprinted on glass coverslips. (a) Stochastic optical reconstruction microscopy (STORM), also known as single-molecule localization microscopy (SMLM), image of cellulose fibril network that was peeled off a protoplast cell—after cell wall regeneration for 48 hours—and adsorbed onto the poly-L-lysine-coated coverslip surface. Cellulose was labeled with Alexa568-tdCBM3a. Alexa568 was induced to blink (see *SI Video 10*) under strong 561 nm excitation and STORM buffer (*SI Materials and Methods*), and single molecules were identified and localized to reconstruct the STORM image. Inset: Comparison of TIRFM (left panel) and STORM (right panel) images of a single cellulose microfibril. (b) TIRFM (left panel) and STORM (right panel) images of closely located multiple fibrils. The cellulose network was heterogenous in the mesh size, with areas of coarse (c) and fine (d) cellulose fibril mesh sizes that were seen to coexist. Various types of inter-fibril associations could be identified from the STORM images: cross-coalescence (arrow); tangential fusion of two curved fibrils (two-headed arrow); and close but non-coalesced two fibrils (chevron). (e) A bundle of two fibrils maintaining a double-stranded fibril segment (∼1.5 μm) in the middle. (f) Scanning electron micrograph (SEM) of a branching cellulose fibril extracted from regenerated protoplast cell walls. Scale bar: 5 μm (a), 0.5 μm (left panel of inset-a), 50 nm (right panel of inset-a), 0.5 μm (b-d) and 0.25 μm (e-f).

## DISCUSSION

Textbook plant cell model schematics typically represent plant primary cell walls as cellulose fibrils tethered by hemicellulose inside an amorphous pectin hydrogel matrix (101). This simple cartoon model, which is based on limited evidence collected over the last hundred years, has been challenged and revised by several experimental/modeling studies, advancing our understanding of how cell walls can give rise to the remarkable mechanical properties critical to support plant growth, development, morphogenesis, and function (5). However, cell wall studies to date have focused mainly on molecular architecture of the existing or already matured cell wall types in various plant tissues and developmental contexts; hence, little is known about the dynamic processes through which the primary cell wall is constructed *de novo* from the cellulose building blocks and developed into the complex ultrastructures as observed by microscopy. In this work, we directly visualized how the Arabidopsis protoplast develops a new cellulose fibril network on the cell surface during cell wall regeneration. We could spatiotemporally resolve the dynamics of this process by directly imaging nascent cellulose biosynthesis, fibrillar growth, and assembly into a fibrillar network over extended time periods (*i.e*., continuously for ∼1 day at 6 min imaging intervals or for several hours at 20 sec imaging intervals) via tdCBM-labeling and TIRFM. Based on our findings, we propose a four-stage model for cellulose fibril network development during cell wall regeneration on plant protoplast surfaces (Fig. 9).

**Figure 9:**
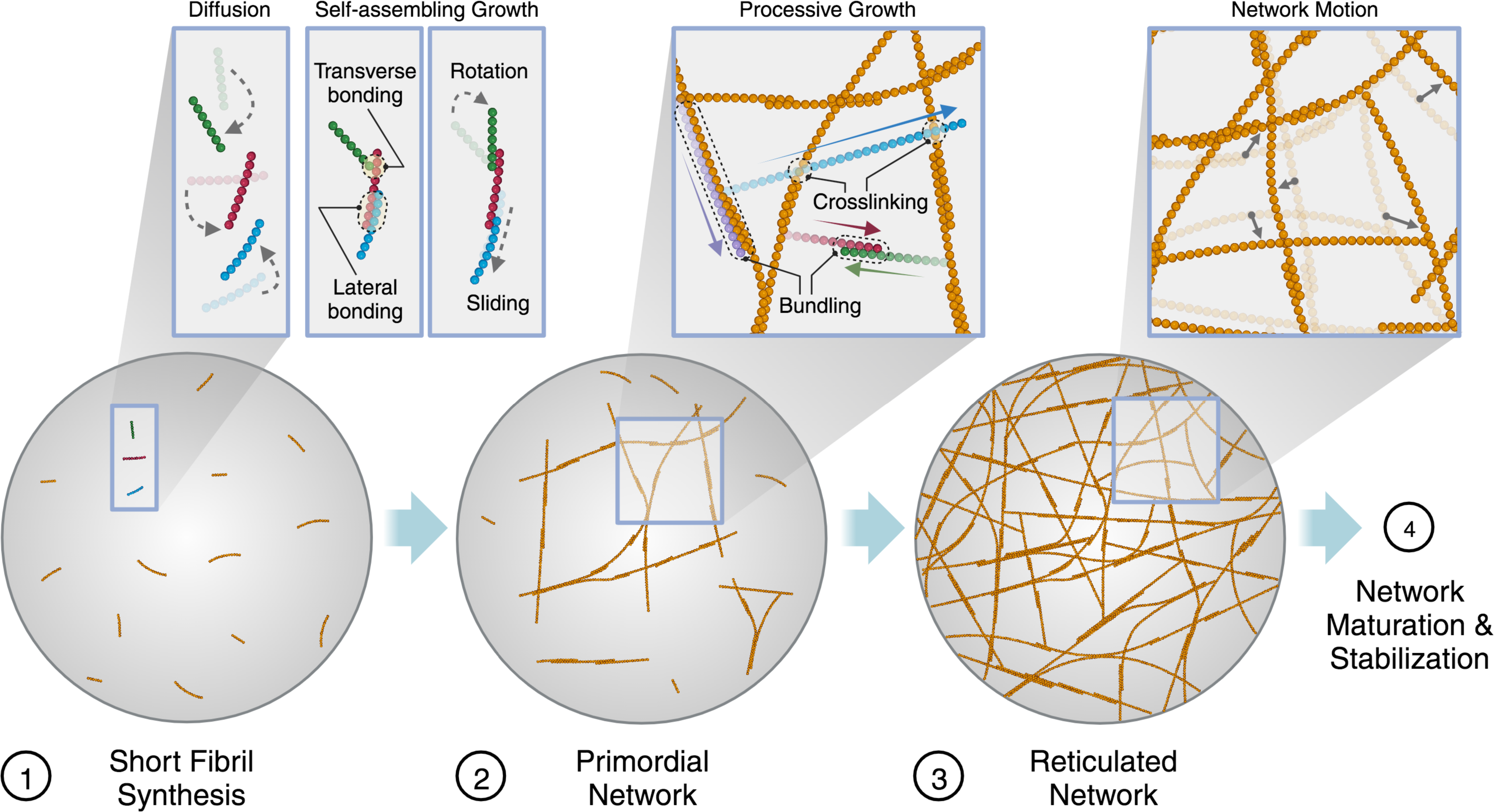
Proposed new model for cellulose biosynthesis and fibril network development during cell wall regeneration in plant protoplasts. Plant protoplasts regenerate the cellulose network during multiple stages of development in this newly proposed model. Cellulose elementary fibrils synthesized by CSC in short lengths dynamically diffuse and coalesce together to form thicker and long fibrils in Stage ①. A less mobile primordial or proto-network emerges from this self-assembly process in Stage ②, which is accompanied by more frequent processive cross-network synthesis events of new single cellulose fibrils, generating a highly reticulated network with denser and finer mesh sizes in Stage ③. The fibrillar network slowly and continuously rearranges during cell wall regeneration, which helps matures its shape to evolve into a more compact/rigid/stable network in Stage ④.

First, nascent cellulose fibrils are synthesized as relatively short fragments (≲ 2 μm) and deposited on the cell membrane surface upon initiation of the cell wall regeneration (Stage ① in Fig. 9). These cellulose fragments randomly diffuse on the surface of the plasma membrane (Fig. 4). At this time, it is not clear whether the observed short fragments are free cellulose fibrils, or they are tethered to the plasma membrane-bound CESA/CSC and undergo active, albeit slow, synthesis/growth while diffusing randomly on the surface. In previous studies, CSCs labeled with fluorescent protein tags were also found to move either along or independently of cortical microtubules in plant tissues (68, 75). The pattern of random translational/rotational motion of the small cellulose fragments in our data seems to be incompatible with the microtubule-guided CSC motion model; it rather implies that either free cellulose is released from CSC to be randomly deposited on the cell membrane surface and/or that a short cellulose fibril is tethered to an inactive CESA/CSC under movement independently of microtubules. Both models could explain observed diffusive motion of short fibrils (Fig. 4), although the origin of this phenomenon needs to be further examined in the context of its correlation with CSC and/or microtubule motions by multi-color fluorescence imaging studies in future work.

Second, the highly mobile cellulose fragments, which are produced in the first stage, frequently encounter each other as they diffuse on the cell surface, and then coalesce together to form thicker and/or longer fibrils (Fig. 5a) that can further self-assemble into a primordial fibril network (i.e., proto-network) with slower mobility and coarse mesh (Stage ② in Fig. 9). Our data suggest that fibril-fibril assembly is predominantly driven by direct cellulose-cellulose interactions between fibrils undergoing rapid diffusion on the cell surface (see further discussion below). Hydrogen bonds, dispersion force, and hydrophobic interactions may be responsible for such cellulose inter-fibril interactions driving the formation of a self-assembled cellulose fibril proto-network (102–104).

Third, processive synthesis of new cellulose fibrils across previously deposited fibrils in the proto-network makes the mesh denser, generating a reticulated network (Stage ③ in Fig. 9 and Fig. 6). Interestingly, processive cellulose fibril elongation is rarely observed for fast diffusing small cellulose fragments at early time periods of regeneration (≲ 10 hour), whereas it occurs more frequently upon emergence of a proto-network, wherein growing cellulose fibrils are mostly tethered to the underlying stationary cellulose network. This biphasic behavior in cellulose synthesis rate is reflected in the temporal change of total Alexa568-tdCBM3a fluorescence seen per single cell in TIRFM data, showing a lag phase of about 10 hours before a fibril network pattern emerges (Fig. 3B and C). Moreover, new fibrils typically grow across the underlying fibrils in the post-lag phase. Such crossover motions are indicative of CSC trajectories generated by the cortical microtubule guidance mechanism (75), although it needs to be further verified by colocalization study of cellulose fibrils and microtubules via two-color fluorescence imaging of live cells in future work. The two types of CSC guidance—autonomous and microtubule guidance systems (75)—may be at play in the biphasic cellulose synthesis behavior observed during cell wall regeneration. It is possible that a CSC actively synthesizing cellulose, either through autonomous or microtubule guidance mechanism, may be susceptible to destabilization, inactivation, and/or disassembly when the nascent cellulose fibril emerges untethered, resulting in highly fragmented cellulose fibril products early in cell wall regeneration. In contrast, CSC may be able to more stably and steadily synthesize longer cellulose fibrils, being guided along a microtubule, when the nascently growing fibril is attached to a semi-stationary proto-network that is constructed by the self-assembly mechanism (Stage ② in Fig. 9). Hence, formation of a primordial cellulose fibril network (*i.e*., proto-network) after the initial lag phase likely provides a crucial template for additional microtubule-guided robust, processive cellulose synthesis by CSC to more effectively develop the final interwoven fibril networks.

Finally, the self-assembled proto-network superstructure is not stationary but rather it is dynamic, capable of slowly and continuously rearranging its shape to become a more compact/rigid/stable network (Stage ④ in Fig. 9 and Fig. 7). Previously published work has identified three distinctive types of cellulose microfibril motions (both experimentally in cell wall stretching assays and computationally in coarse-grained molecular simulations)—sliding, straightening, and changes in bundling. These distinct types of motions were proposed to explain the mechanical response of the cell wall to external forces (78, 98). They are also likely to be involved in the cellulose network dynamics observed in our study, possibly playing important roles in relieving any local buildup of mechanical stress that may occur while nascent cellulose is continually deposited on the protoplast surface during cell wall development.

A model of the primary cell wall as a collection of well-separated cellulose microfibrils mechanically tethered by xyloglucan polymer bridges (101) is widely accepted. However, whether xyloglucan plays a major load-bearing role has been called into question by studies of xyloglucan-deficient Arabidopsis double mutants (*xxt1, xxt2*), which showed only minor effects either on growth and appearance of the mutant plant (90) or on cell wall regeneration of the mutant protoplast (81). An alternative model postulates that cellulose microfibrils make tight contacts between each other to form cellulose-cellulose junctions, termed “biomechanical hotspots”, that play key structural and mechanical roles in formation and stabilization of the primary cell wall (5, 105). Such junctions could originate from either direct inter-cellulose interactions or mediation, at least in part, by xyloglucan. Our time-lapse TIRFM data (Fig. 3, 5, and 6) and STORM images (Fig. 8) confirm the existence of junctions between cellulose fibrils; moreover, they uncovered the dynamic processes by which such junctions form through fibril coalescence, yielding stable cellulose fibril networks on the cell surface. To investigate the role of xyloglucan in cellulose fibril coalescence and cell wall development, we acquired time-lapse TIRFM images of cellulose in xyloglucan-deficient Arabidopsis double mutants (*xxt1, xxt2*)-derived protoplasts during cell wall regeneration (Fig. S6, *SI Video 11*). Mutant protoplasts showed cell wall regeneration behaviors very similar to wild-type, suggesting the primary role of direct cellulose-cellulose interaction in the formation of inter-fibril junctions despite the absence of xyloglucan.

Protoplasts have been employed as a powerful tool for nearly six decades to study various molecular and cellular processes such as cell wall synthesis, regeneration and modulation against intrinsic and extrinsic stress factors (106, 107). However, removal of the cell wall is known to activate stress/wounding response and metabolic process function genes (108, 109), and this can be a major limitation of using isolated mesophyll protoplasts as a proxy to understand the mechanism of cell wall biosynthesis as compared to intact or natural plant tissues undergoing cell wall growth. It is thus possible that the mechanism of cell wall biosynthesis during normal plant cell development could be distinct from that observed using isolated protoplast cells. Nevertheless, protoplasts competent for cell wall regeneration provide a unique opportunity at this time to enable the study of *de novo* cell wall biogenesis, with minimal background from a pre-existing cell wall matrix. This system makes possible the stepwise description of the dynamic processes that facilitate evolution and self-assembly of the intricate cell wall, starting with short, highly diffusive cellulose fragments. Whether the processes that we described here for regenerating protoplast also take place during some context of normal development will have to await new techniques that could overcome the background issues posed by pre-existing cell wall materials.

In summary, we developed a live-cell imaging system to observe cellulose formation and fibrillar assembly on the surface of plant protoplasts undergoing cell wall regeneration. Combining fluorophore labeled tandem CBM3a, TIRFM imaging, and ambient light/temperature control enabled us to continuously monitor the striking dynamics of nascent cellulose microfibril synthesis and development toward a network during the initial ∼1 day of cell wall regeneration in high spatial and temporal resolution. The novel findings captured by our study provide important insights into dynamic aspects of plant cell wall and cellulose fibril synthesis at the single-cell level. Our microscopy platform and experimental workflow will also be suitable for studying the correlative dynamics of other cell wall components going beyond the role played by cellulose in live plant protoplasts.

## ACKNOWLEDGMENTS

This work was primarily supported by the Department of Energy (DOE) award DE-SC0019313. S.-H.L. and S.P.S.C. also acknowledge partial support from the Rutgers Office of Research & Innovation. S.P.S.C. acknowledges partial support from his NSF CBET Career Award (1846797). S.-H.L acknowledges partial support from Rutgers New Faculty Startup Fund through the Institute for Quantitative Biomedicine. We thank Ryan Johnson and Lee Alexander from the GLBRC biomass analytical facility for conducting the composition analysis on protoplast samples. The GLBRC core facility work is supported in part by the Great Lakes Bioenergy Research Center, U.S. Department of Energy, Office of Science, Office of Biological and Environmental Research under Award Number DE-SC0018409. We thank Stephen Burley (Rutgers University) and Kenneth Keegstra (Michigan State University) for providing helpful comments on the manuscript.

## AUTHOR CONTRIBUTIONS

S.P.S.C., E.L., and S.-H.L. conceived and supervised the study. H.H. led and conducted all the imaging experiments including SEM, confocal, TIRFM, and STORM microscopy work. D.J. developed, characterized, and produced all CBM-based dyes used for imaging. D.J. led all biochemical and cell wall composition experiments and data analysis. H.H., M.I., D.J. and J.S. developed and validated the plant protoplast cell wall regeneration assay. J.S. maintained the Arabidopsis plants and generated protoplasts for the team. H.H. and S.-H.L. analyzed all the microscopy data. D.J., with the assistance of H.H., prepared an initial incomplete manuscript draft along with critical inputs from S.P.S.C., E.L., and S.-H.L.. S.-H.L. prepared the final figures/videos and wrote the final complete version of paper. All the authors reviewed, edited, and approved of the final manuscript draft.

## Supplementary Information (SI) Appendix

**Supplementary Information Table of Contents**

Supplementary Videos S1-S11 (included as separate SI media files with their descriptions in SI appendix pdf; pages 2-3)

Supplementary Figures S1-S6 (included in SI appendix pdf; pages 4-9)

Supplementary Materials and Methods (included in SI appendix pdf; pages 10-16)

Supplementary References (included in SI appendix pdf; pages 17-18)

### Supplementary Videos Description

**Video 1 [supplementary to Fig. 2a].** Time-lapse TIRFM image sequence of cellulose fibrillar network being continuously degraded by a commercial cellulase (Cellic CTec2, Novozymes). Cellulose was regenerated on a protoplast cell surface for 24 hours and labeled with Alexta568-tdCBM3a. Acquisition speed: 1 frame per 2 minutes.

**Video 2 [supplementary to Fig. 3b].** Time-lapse TIRFM image sequence of cellulose synthesis and fibril formation on a protoplast surface in a regeneration media (WI/M2). Cellulose was in situ stained with Alexa568-tdCBM3a. Image acquisition began 1 hour after the start of cell wall regeneration and continued for ∼16 hours. Acquisition speed: 1 frame per 6 minutes.

**Video 3 [supplementary to Fig. 4a].** Time-lapse TIRFM image sequence demonstrating random diffusive motion of short cellulose fibril fragments nascently synthesized and deposited on the surface of protoplast at the early stage of cell wall regeneration. The trajectory of a cellulose fragment is overlaid for better visualization of the motion. Cellulose was in situ stained with Alexa568-tdCBM3a. Acquisition speed: 1 frame per 2 minutes. Scale bar: 2 μm.

**Video 4 [supplementary to Fig. 5a].** Time-lapse TIRFM image sequence showing coalescence events between nascently synthesized cellulose fibrils on the protoplast surface, which lead to generation of thicker and/or longer fibrils. Acquisition speed: 1 frame per 20 seconds. Scale bar: 2 μm.

**Video 5 [supplementary to Fig. 5b].** Time-lapse TIRFM image sequence showing dynamic morphological changes of a coalesced cellulose fibril on the protoplast surface. Acquisition speed: 1 frame per 20 seconds.

**Video 6 [supplementary to Fig. 5c].** Time-lapse TIRFM image sequence showing dynamic bending and unbending of an extended cellulose fibril on the protoplast surface. Acquisition speed: 1 frame per 2 minutes.

**Video 7 [supplementary to Fig. 5c].** Time-lapse TIRFM image sequence showing an extreme bending (nearly 180°) event that leads to an intra-fibrillar coalescence of a single cellulose fibril on the protoplast surface. Acquisition speed: 1 frame per 2 minutes.

**Video 8 [supplementary to Fig. 6a].** Time-lapse TIRFM image sequence showing processive growth of single cellulose fibrils on the protoplast surface. Acquisition speed: 1 frame per 2 minutes.

**Video 9 [supplementary to Fig. 7a].** Time-lapse TIRFM image sequence showing a cellulose fibril network whose morphology slowly but continuously changes on the protoplast surface as the cell wall regeneration process goes to completion. Left panel: raw images. Right panel: images processed with Laplacian of Gaussian filtering. Acquisition speed: 1 frame per 20 seconds.

**Video 10 [supplementary to Fig. 8a].** Time-lapse TIRFM image sequence showing fluorescence blinking of Alexa568-tdCBM3a dye that labels a surface-adsorbed protoplast cellulose fibril network. The fluorescence blinking events, which was induced under a strong 561nm laser excitation and a special buffer condition, are analyzed to produce a super-resolution image (Fig. 8a). Acquisition speed: 33 frames per second. Scale bar: 5 μm.

**Video 11 [supplementary to Fig. S8a].** Time-lapse TIRFM image sequence of cellulose synthesis and fibril formation on the surface of a xyloglucan knockout protoplast—(xxt1, xxt2) mutant—in a regeneration media (WI/M2). Image acquisition began 2 hours after the start of cell wall regeneration and continued for ∼24 hours. Acquisition speed: 1 frame per 6 minutes.

## Supplementary Methods

### Plant Materials and Growth Conditions

*Arabidopsis thaliana* Col-0 (Columbia) was used for all wild-type or transgenic control experiments. Seeds of Col-0 were purchased from ABRC (Arabidopsis Biological Resource Center). Approximately 50 seeds were incubated in 10% bleach for 5 mins, and washed thrice with excess sterile distilled (DI) water, and resuspended in 50 µl of DI water before germination. Sterilized seeds were poured onto Murashige and Skoog (MS) media plates supplemented with Gamborg’s vitamins (Phytotech lab, M404). Seeds were spread evenly using a sterile pipette tip and incubated at 4 °C and ambient humidity for maximum 3 days. Plates were later transferred to the plant growth chamber at 23 °C, with a light intensity of 120 - 150 µmol/m^2^/s with 40–65% relative humidity and a 10-hour photoperiod consisting of 10 hours of light and 14 hours of dark photoperiod. Seedlings were allowed to grow for 8-10 days before transferring into pots with promix soil (Grinffin, BK25-V) and incubated in the growth chamber with the same conditions. All other reagents were purchased from VWR, Thermo Fisher Scientific, and Sigma Aldrich unless mentioned otherwise.

### Arabidopsis Mesophyll Protoplast Isolation

Mesophyll protoplasts were isolated, as described by Yoo et al. (1), with slight modifications as described here. Briefly, 3-4 weeks old fully expanded leaves of Arabidopsis plants (as shown in Figure 2a) were selected. The selected leaves were uniformly sized to avoid heterogeneity in the size of protoplasts. 0.5-1 mm thick leaf strips were cut off from the middle part of a leaf using sharp and sterile razor blades(single-edged blade; VWR Scientific, cat. no. 55411-055) without any damage to the tissue at the cutting site. The leaf strips were quickly transferred into a sterilized enzymatic solution (0.4 M Mannitol, 20 mM KCl, 20 mM MES, 1.5% Cellulase R10 (Yakult Pharmaceutical Ind. Co., Ltd., Japan), 0.4% Macerozyme R10 (Yakult Pharmaceutical Ind. Co., Ltd., Japan), 10 mM CaCl2, 5 mM 2-mercaptoethanol, 0.1% BSA) using flat-tip forceps and submerged completely. The leaf strips were vacuum infiltrated for 30 minutes before stagnantly incubating the samples in the dark for maximum 5 hours for enzymatic digestion. Upon incubation, the protoplasts were diluted with an equal volume of sterile W5 solution (154 mM NaCl, 125 mM CaCl2, 5 mM KCl, 2 mM MES in double-distilled water). The undigested leaf tissues were removed by filtering the solution through a pre-washed nylon mesh (75 µm, laboratory sifters, Carolina Biological Supplies, cat. no. 65-2222N). Protoplasts were collected by spinning at 207 rcf for 3 minutes at 4 °C. The isolated protoplasts were washed twice with 10 ml of W5 solution to remove any residual enzyme solution. Finally, the protoplasts were resuspended in 0.5 ml of W5 solution. The quality of the protoplasts isolated was routinely observed under the regular light microscope (Fig. 1). The quality of protoplasts was deemed unsuitable if they were too large (>100 microns), too small, or broken/lysed. As a rule of thumb, medium-sized leaves were chosen for isolation to generate protoplasts with overall size in the rage of 30-50 μm under bright-field microscopy. The concentration of protoplasts was measured using a hemocytometer (Hausser Scientific, cat. no. 1483) and the final concentration was adjusted to 2 x 10^5^ protoplasts ml^-1^ of W5 solution. The freshly isolated protoplasts were stored on ice for immediate use for experiments typically within 2 hours, or in a 4°C fridge for experiments in the following day.

### Protoplasts Cell Wall Regeneration on Lab Bench

Approximately 200 µl of protoplasts of concentration 2 x 10^5^ protoplasts ml^-1^ were used for the cell wall regeneration experiments. The protoplasts were gently pelleted down at 207 rcf for 2 minutes using a refrigerated centrifuge (4°C). The supernatant was removed, and the protoplasts were then stored in a WI solution media solution (4 mM MES, 0.5 M mannitol, 20 mM KCl). Cell wall regeneration media (WI/M2) contains an equal volume of WI and M2 media (Gamborg’s B-5 basal medium with minimal organics 6.4 g/L from Sigma, 0.8 M trehalose, 0.1 M glucose, 1 µM 1-naphthalene acetic acid, pH 5.7) (2). The protoplast suspension in cell wall regeneration media was incubated at room temperature (20 - 23 °C) under continuous ambient light exposure conditions (Philips hue lamp) over 24 hours.

### Protoplast Samples Preparation for Cell Wall Glycans Composition Analysis

Approximately 1.5 mg of air-dried (18 hours in oven at 56 °C) protoplasts were transferred into 2-mL reaction tube (Sarstedt). To remove the cell wall regeneration media components, the samples were washed with 1 mL of double-distilled water and vortexed for 15 seconds at maximum speed (>13,000-g) in a mini-centrifuge. The aqueous phase is collected at each step for analysis. The protoplasts were then recovered after washing by centrifuging into a pellet at 10,600 x g for 30 seconds. The supernatant was removed carefully without disturbing the solid pellet. Samples were then washed with 1 mL each of 70 v/v% ethanol followed by chloroform:methanol (1:1 v/v) mixture in series. Following that, the sample was resuspended and washed with 1 mL double-distilled water twice. All aqueous phase fractions were pooled together and filtered using 0.2 µm nylon syringe filters. 1 mL of the aqueous extract was evaporated in vacufuge and lyophilized overnight. Alternatively, the washed protoplasts were also placed inside a lyophilizer (Labconco FreeZone 4.5) for drying overnight. Finally, the samples were washed with 1 mL acetone to collect the residual pellet samples stuck to the walls of the tube and dried overnight. The dried solid samples were stored at 22 °C for cell wall polysaccharides analysis.

### Trifluoroacetic acid (TFA) Hydrolysis and Alditol Acetate Composition Analysis

To analyze non-crystalline or amorphous cellulose and other matrix polysaccharides, the dried/washed samples were initially subjected to mild TFA hydrolysis. Here, 250 µl of TFA solution (14.95 mL of 2 M TFA and 1.3 mL of 5 mg/mL inositol stock solution) was gently added to the walls of each sample tube. Samples were later incubated for 1.5 hours at 121 °C on dry bath and the heating block then cooled down on ice. The insoluble residue after TFA hydrolysis was pelleted out by centrifuging at 10,000 rpm for 10 minutes. Next, 25 µl of the solubilized TFA hydrolyzate supernatant was transferred into three separate tubes each without disturbing the solid pellet. Then, 200 µl of isopropyl alcohol (IPA) was added to one of the TFA hydrolyzate glass tubes and dried at 25 °C under moderate airflow (∼2 psi) for 35-40 minutes. The resulting small salty pellet was washed with IPA three more times (for a total of four additions) and blow dried at 25 °C. The dried tubes were stored inside a dry fridge overnight at 22 °C. To the dried TFA hydrolysate, 100 µL of fresh sodium borohydride solution (10 mg sodium borohydride in 10 mL of 1M ammonium hydroxide) was added and vortexed at maximum speed. The suspension was incubated at room temperature for 1.5 hours. The sample was neutralized using 150 µl of glacial acetic acid. In the next step, 150 µl of glacial acetic acid and methanol mixture (as a volume ratio, 1:9) was added and evaporated at 35 °C under moderate airflow (∼2 psi) for 35-40 minutes. This step was repeated, and the resulting white crystalline residue was resuspended in 250 µL of methanol (MeOH) and evaporated at 35 °C twice. To the opaque white residue (dried alditol precipitate), 50 µL of pyridine and 50 µL of acetic anhydride was added and vortexed thoroughly. Samples were incubated for 20 minutes at 121 °C before cooling the heating blocks on ice. Next, 200 µL of toluene was added and evaporated at 25 °C under a gentle airflow (∼1 psi) for 10-15 minutes. The addition of toluene and evaporation step were repeated to obtain a white salty residue. Since the alditol acetate derivatives are volatile, the evaporation processes did not take more than 10 minutes and were instead stored in a 22 °C dry fridge overnight when needed. To the dried alditol acetate derivatives, 1 mL of ethyl acetate (EtOAc) and 1 mL of double-distilled water were added in series and vortexed thoroughly. The phases were separated by centrifuging the tubes at 2,000 rpm for 1 minute. Finally, 1.75 mL of double-distilled water was added to increase the volume for pipetting EtOAc from the top phase and 200 µL of EtOAc layer was transferred into glass GC vials (with glass inserts), and the samples were stored at 4 °C overnight or analyzed using GC-MS (Agilent 7890A) directly to estimate the TFA acid hydrolysis released monosaccharides present (as monosaccharide equivalents on dry weight basis of original dried protoplast samples).

### Updegraff Method for Estimating Mild Acid Resistant Crystalline Cellulose

To analyze the crystalline cellulose present in regenerated protoplasts analyzed as dried solid samples, 1.5 mL of Updegraff reagent (acetic acid: nitric acid: water = 8:1:2, volume ratio) was added to the TFA hydrolysate and vortexed thoroughly. The samples were heated at 100 °C for 30 minutes, and the solid material was recovered, as detailed in the previous section. The samples were resuspended in 1.5 mL of double-distilled water and 1.5 mL of acetone twice following centrifugation and recovery of cell pellet after each step. Samples were dried at room temperature overnight. Next, 150 µL of 72% sulfuric acid was added to the dried sample and incubated at room temperature for 30 minutes. After incubation, 875 µL of double-distilled water was added to the samples and vortexed thoroughly. Tubes were centrifuged at 10,000 rpm for 5 minutes, and the supernatant was collected. 10 µL of the supernatant was mixed thoroughly with 90 µL of double-distilled water and 200 µL of freshly prepared 2 mg/ml anthrone solution (anthrone in sulfuric acid) in a 96-well plate. The plate was incubated on a plate heater at 80 °C for 30 mins. The incubated plate was later equilibrated to room temperature for 30 mins before measuring the absorbance at 405 nm using a plate reader (Spectramax Plus 384). Samples were compared to the glucose standards, and the error bars are the standard deviations of the mean. All experiments were performed with at least two replicates.

### Enzymatic Hydrolysis of Protoplast Cell Walls by Cellulases

Fixed protoplasts after cell wall regeneration were used to examine the enzymatic deconstruction in the presence of commercial crude cellulolytic cocktails or purified cellulases. The fixed protoplasts were isolated from the 12% sorbitol and were washed with 25 mM sodium acetate buffer (pH 4.5) twice. The washed protoplasts were incubated with 1 µg/µl of commercially available cellulase cocktail (Cellic CTec2, Novozymes) at room temperature for live imaging of cellulose fibril degradation visualization. See below for live cell imaging protocol. The supernatants with released monosaccharides were collected and stored at −20 °C for reducing sugars analysis, if needed. Similarly, to examine the selective degradation of cellulose and xyloglucan in the regenerated protoplasts, purified glycosyl hydrolases (E-CELBA from Neogen Corp.) specific towards either cellulose and xyloglucan were used. The fixed protoplasts were incubated at 40 °C for 2 hours with the enzymes prior to imaging. The supernatants were collected and subjected to DNS reducing sugars assay, as reported previously (3).

### Preparation of CBM Probes for Live Cell Imaging

It was imperative to use a dye or fluorophore-labeled protein probe in the regeneration media that was stable over extended time durations, had minimal cell toxicity, and did not significantly interfere with the growth and assembly of plant cell walls. In addition, the probe had to be highly specific and largely irreversible (i.e., with high association rate but low dissociation rate) in its binding properties to capture even nascent cellulose chains emerging from the plasma membrane. Tandem carbohydrate-binding modules (CBMs) or repetitive CBMs exist in nature alongside fungal cellulases and provide high binding specificity to various substrates (4). Depending on the type of CBM and the matrix polysaccharide to be imaged, different CBMs have been utilized as an alternative to classical immunolabeling methods to localize plant cell wall glycans (5). CBM3a is widely preferred due to its ability to recognize semi-crystalline cellulose in plant cell walls and tissues. Here, we conjugated the Alexa-Fluor 568 dye to a novel tandem CBM3a construct that bound to cellulose. The succinimidyl ester of Alexa-Fluor dye is an amine-reactive species and quickly conjugates with the primary amines (of lysine) and amine terminus of proteins. In our previous study, we observed that tandem CBM3a has a higher binding rate than the single CBM3a and could be readily used for real-time imaging of polysaccharides biosynthesis in plants (6).

Expression and purification of tandem CBMs was performed as described previously (6). pEC-GFP-CBM3a is an *E. coli* expression vector and was kindly provided by the Fox lab (UW Madison) (7). Other CBMs (i.e., CBM17 and CBM1) were prepared as reported elsewhere (8). The plasmid was also used as a backbone to insert CBM3a (*Ruminoclostridium thermocellum*) or other CBM genes through the sequence and ligation-independent cloning (9, 10). Sequence verified plasmid DNA was expressed in *E. coli* BL21-CodonPlus-RIPL [λDE3] and purified using immobilized metal affinity chromatography (IMAC) as mentioned previously (6, 11). For labeling, the tandem CBM3a in PBS buffer was incubated with the fluorophore (Alexa Fluor 568 NHS Ester) at a molar ratio of 1:10 at 4°C and kept in darkness for overnight. The excess fluorophore was removed using gel filtration spin columns (Bio-Rad) according to the manufacturer’s specifications. Labeled samples were aliquoted and flash frozen using liquid nitrogen before storing at −80°C.

### Confocal Microscopy

We fixed the sample with 0.001% glutaraldehyde and added glycine solution (final concentration is 5 mM) as a quenching agent. The fixed protoplasts were incubated in the presence of 200 nM fluorescent dye-labeled tandem CBM3a and washed out the unbound probes with PBS buffer. Secondly, LM15 antibody was added to the sample at a dilution of 1:20 in PBS buffer. After three washes with PBS buffer, an anti-rat IgG Alexa-Fluor secondary antibody was added at a 1:2,000 dilution and the samples were incubated for an additional 1 hour. The samples were stained with CBM probes (or calcofluor white or LM15 antibody) without washing step and 20 μL of the sample was injected into the sandwich-type chamber assembled with 25 mm round coverslip, GeneFrame (AB0576, Thermo Fisher Scientific) and slide glass (470150-480, VWR International). Confocal imaging was performed with a commercial confocal microscope (Leica TCS SP8, Leica Microsystems) equipped with a 63x/1.30 glycerin objective lens and two laser lines, 488 nm and 561 nm) and the stacked image data were constructed by using ImageJ software (NIH) (12).

### Live Plant Cell Imaging via Total Internal Fluorescence Microscopy

We prepared a microwell chamber by attaching a 250 μm-holed silicon spacer (MMA-0250-100-08-01, Microsurfaces Pty Ltd) onto a plasma cleaned 25 mm coverslip (CS-25R15, Warner Instruments) and inserting the coverslip to magnetic chamber (Chamlide-CMB, Live Cell Instrument). We filled the chamber with 200 μL of the protoplasts incubation media (60 v/v% of WI + 40v/v% of M2 along with 0.02% BSA and 100 nM of Alexa568-CBM3a probe) in first and then slowly injected 100 μL of protoplast samples into the same media. The samples were usually preincubated for 2 hours to allow the protoplasts to settle to top of the coverslip. For in situ enzymatic reactions, we replaced the sample buffer by gently washing the regenerated protoplast samples with the imaging media (150 nM of Alexa568-CBM3a and 0.02% BSA in pH 5.5 MES buffer) and left the cell wall containing matured cells for 30 minutes prior to addition of the enzymes. Prior to the image acquisition, we lastly add a cellulase, Cellic CTec2 (0.5 mg/ml, Novozymes) and the total volume did not exceed 400 μL.

For the time-lapse imaging to visualize cellulose or cell wall biosynthesis, we use a homebuilt multicolor TIRF (Total internal reflection fluorescence) microscope (13) with a programmable lamp (Hue lamp, Philips) standing on the upper side of the sample chamber. The lamp is controlled by open-source program (LabVIEW MakerHub, National Instruments) linked to Nikon imaging software (NIS-Elements, Nikon). We set the color of the bulb to warm yellow light: 230 (red), 200 (green) and 100 (blue). And we generated the trigger signal on the NIS-Elements to repetitively switch the lamp off prior to the acquisition. The images were taken constant rate of 20 seconds for the drift motion analysis or few minutes for the long-term imaging over several hours. We typically used 2 mW of 561 nm laser excitation for observing the cell wall synthesis and occasionally did dual-channel imaging with 5 mW of 488 nm excitation for the transgenic line sample prepared from GFP-TUB6 (CS6550, ABRC).

### Normalized Cross-Correlation (NCC) Image Analysis to Quantify Cellulose Mobility

The normalized cross-correlation matrix *ncc*(***A*, *B***) between matrices ***A*** and ***B***, representing two images with pixel values *A_ij_* and *B_ij_*, respectively, is given by

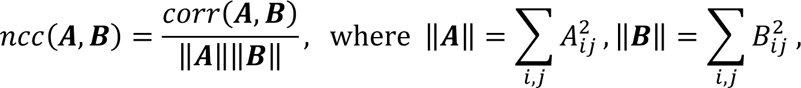

and

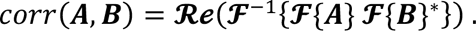

Here, ***F***{} and ***F***^-1^{} represent 2D Fast Fourier Transform (FFT) and inverse 2D FFT, respectively; ***Re***() represents the real parts of complex values; and ()^∗^ represents complex conjugate operation. *ncc*(***A*, *B***) is a matrix (i.e., 2D gray scale image with pixel values between 0 and 1): its center pixel representing the cross-correlation of ***A*** and ***B*** without any shift between the two images, and (*i*, *j*)*^th^* pixel relative to the center representing the cross-correlation calculated with shifting ***B*** from ***A*** by *i*-columns and *j*-rows. When the two input images have no correlation, all the NCC matrix elements will be nearly zero. When two input images are identical, the NCC matrix will be a 2D bell-shaped curve with the peak value of 1 at the center pixel. If the two input images are identical but one is shifted from the other in (x, y) plane, the NCC matrix will be a similar 2D bell-shaped curve with just its peak location displaced from the center pixel by the same image shift amount. Therefore, NCC matrix is commonly used to estimate drift motion with high precision in various microscopy (14, 15).

A fluoresce micrograph typically contains a noise component and also a non-zero, non-uniform background signal, all of which undesirably contribute to NCC. We therefore applied a bandpass filter (16) to the input images to remove both the noise and background signals when calculating NCC. We applied NCC to the time-lapse image sequences of fluorescence-labeled cellulose fibrils on protoplast surfaces to quantify the overall cellulose fibrillar motion by the cross-correlation between two consecutive image frames. The cellulose motion between two consecutive image frames largely originates from two sources: first, common drift either due to motion of an entire protoplast cell or microscope instrumental drift; and second, motion of cellulose fibrils themselves on the protoplast surface. These two types of motion will affect the NCC matrix differently. The drift motion will largely shift the peak position whereas the cellulose motion on the protoplast cell surface will reduce the peak value in the NCC matrix. Therefore, we fitted the NCC matrix with 2D Gaussian functions for a series of two consecutive frames over time; and plotted the peak values in time to estimate the drift-corrected overall cellulose fibril motion on the cell surface (e.g., Fig. 3d and 7c).

### Single Molecule Localization Microscopy (SMLM) of Cellulose Fibril Network

Protoplast cells derived from prc1-1 transgenic Arabidopsis plant line were incubated in the cell wall regeneration media (WI/M2) for 48 hours until cell surfaces were densely coated with cellulose fibrils. The mature protoplast cells were injected onto the top surface of a coverslip coated with poly-L-lysine (P4707, Sigma Aldrich). We manually shook the coverslip to carefully peel off the cell wall from the plasma membrane, and then removed the media and other debris by pipetting and washing with PBS buffer (pH 7.4). We mounted the coverslip onto a magnetic chamber and stained the sample with Alexa568-CBM3a. For the photo-switching, the PBS buffer was replaced by a fresh imaging buffer mixed with 1xPBS pH7.4, glucose oxidase (G2133, Sigma-Aldrich), catalase (C9322, Sigma-Aldrich), Trolox (238813, Sigma-Aldrich) and BME (63689, Sigma-Aldrich). We acquired 20,000 frames at 33 frames per second through a homebuilt TIRF microscope and the super-resolution image was reconstructed with custom MATLAB codes. The details of the STORM protocol and data analysis are described elsewhere in the literature (17).

### Cellulose Fibril Particle Diffusion Tracking Analysis

We chose several micrometer-sized small fibers from the flat area of the oval-shaped cells and manually tracked each particle using an ImageJ plugin. The speed of drift motion was calculated as the net movement over total tracking time and averaged all the values. For mean square displacement (MSD) analysis, data were analyzed with a custom MATLAB code, which extracted the MSD using a function,< [/(F + τ) − /(F)]^6^ >, for each individual trajectory. The symbol, τ is time delay. Sub-diffusion was analyzed by fitting a linear relation between the log scale of MSD and the log scale of time delay at short time lag (10 frames, i.e., 200 seconds). We determined the anomalous factor from the slope of fitting parameter and graphed the distribution for 23 distinct fibers (Figure 4).

## Supplementary Figures

**Figure S1.**
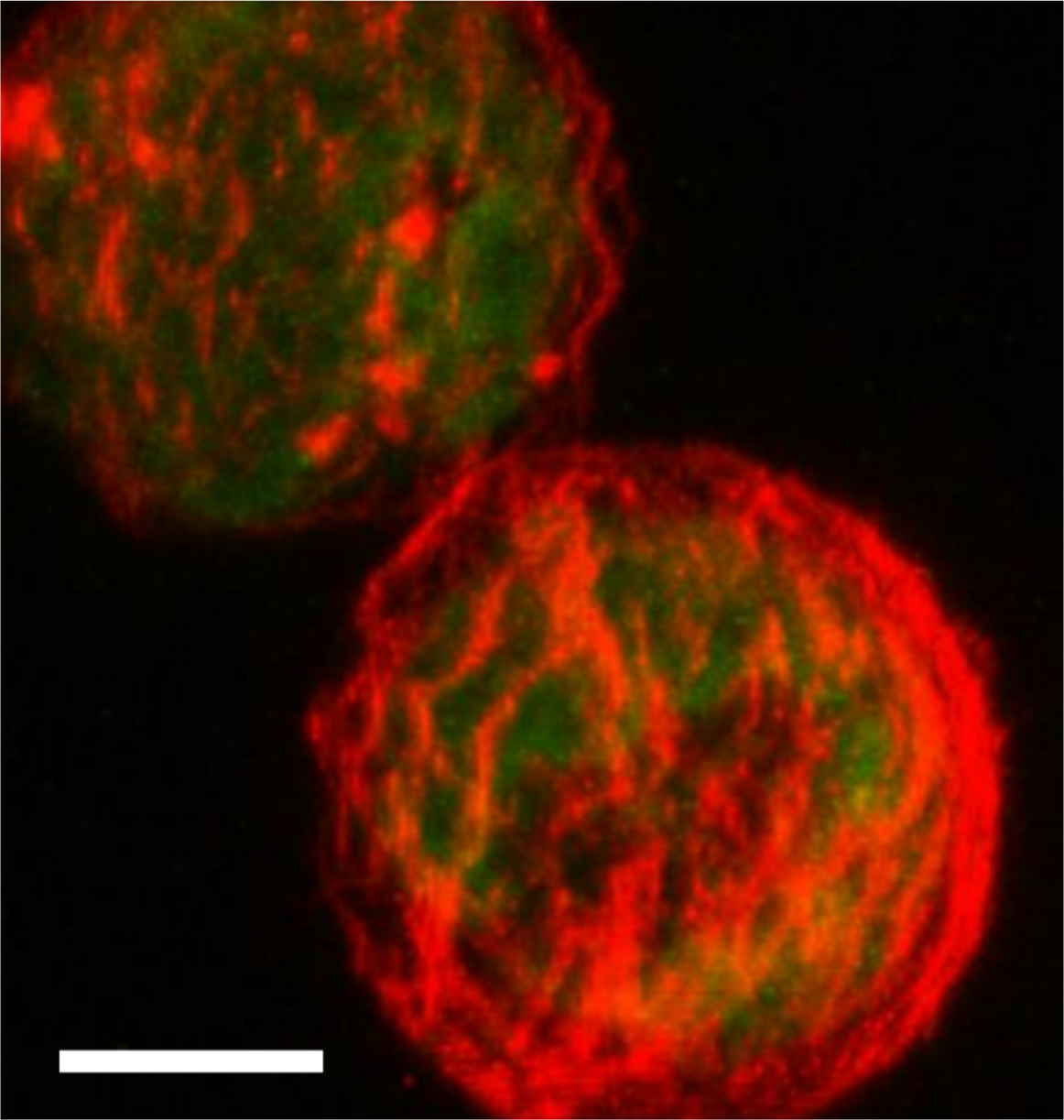
Calcofluor labeling confirms presence of cellulose in regenerated cell walls. A representative confocal image (maximum z-stack projection) of Arabidopsis protoplasts depicting regenerated cell walls after incubation in WI/M2 regeneration media for 18 hours is shown here. Samples were fixed prior to staining with Calcofluor White (CFW) to visualize β-glucans like cellulose present in the regenerated cell wall. Red: CFW fluorescence by 405 nm excitation. Green: Chloroplast autofluorescence by 488 nm excitation. Scale bar: 10 μm.

**Figure S2.**
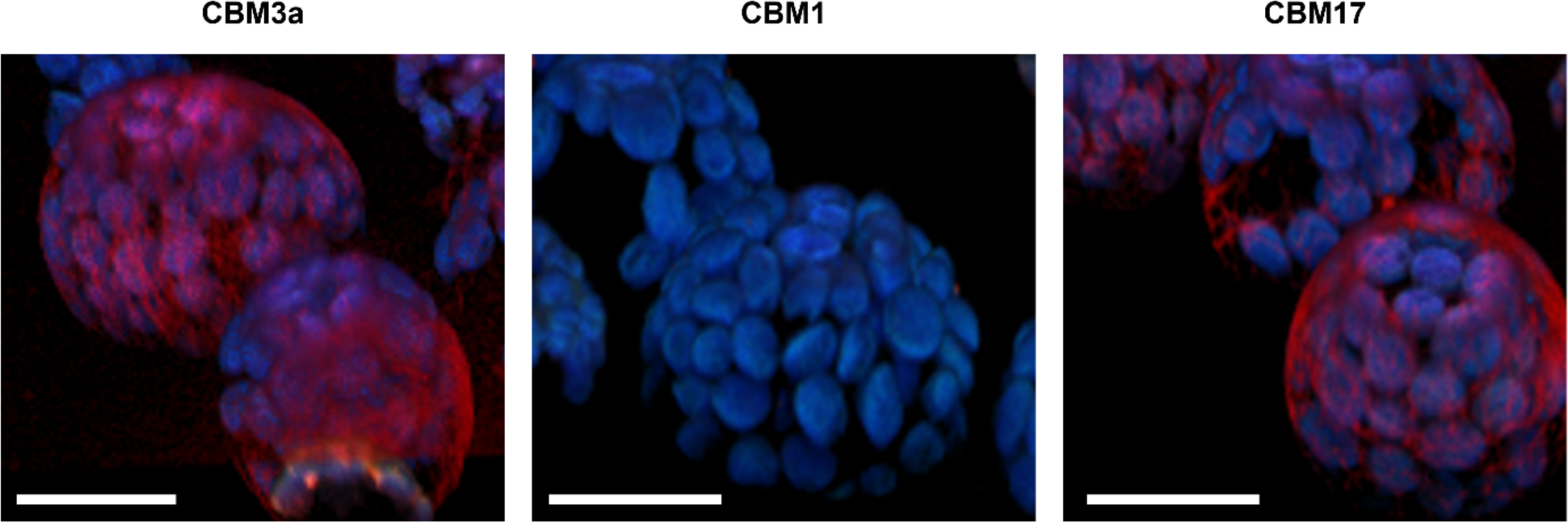
CBM labeling suggests regenerated cell walls are abundant in amorphous or disordered cellulose fibrils. Representative confocal z-stack images of Arabidopsis protoplast cells incubated for 25 hours in the cell wall regeneration media (WI/M2) are shown here. The samples were fixed after cell wall regeneration period and stained with three different Alexa568-CBM probes: CBM3a (left), CBM1 (middle) and CBM17 (right). Except the CBM1-stained cell samples, CBM3a and CBM17 could readily stain the nascent cellulose fibrils covering the plant protoplast cell surface. Red: Alexa568 fluorescence by 561 nm excitation. Blue: Chloroplast autofluorescence by 488 nm excitation. Scale bar: 20 μm.

**Figure S3.**
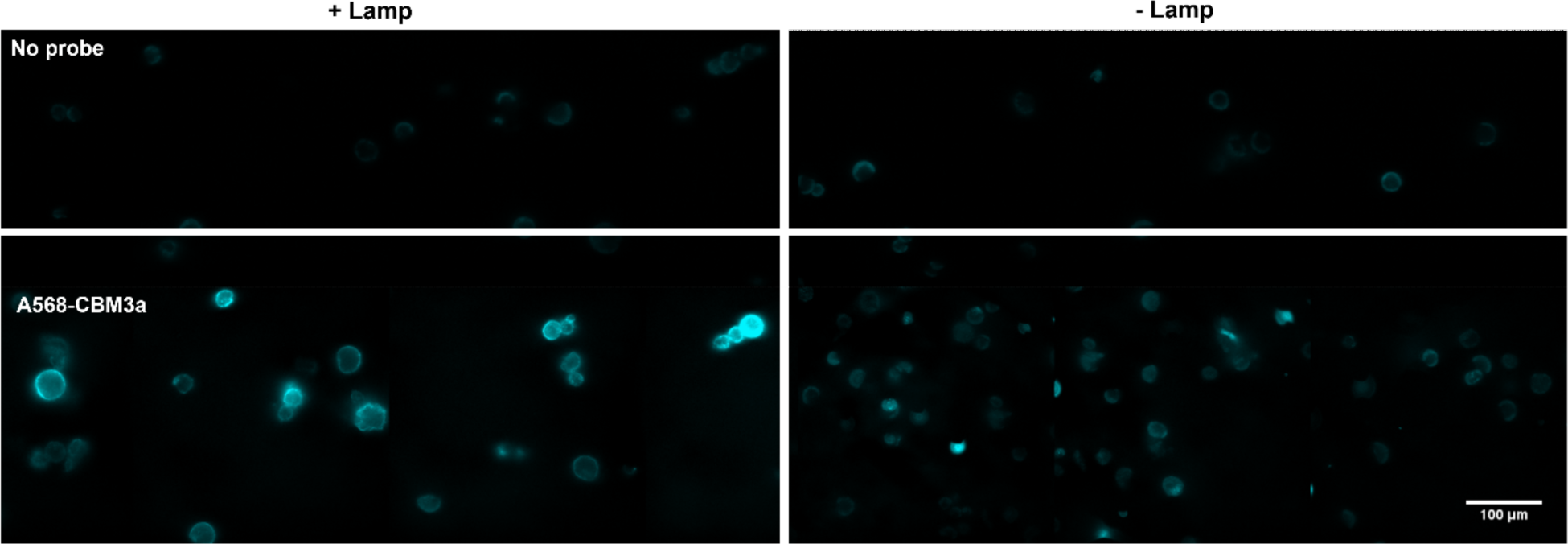
Ambient/white light is critical to protoplasts cell wall regeneration. Epifluorescence images of regenerated Arabidopsis protoplast cell walls (after 18-hour incubation in WI/M2 media) with (left panel) or without (right panel) ambient white light provided by a LED lamp (Hue bulb, Phillips) during the entire cell wall regeneration period. Cell walls were visualized by staining synthesized cellulose with Alexa568-tdCBM3a (561 nm excitation). Ambient light was clearly essential for the protoplasts to reproducibly regenerate cell walls with fibrillar cellulose synthesis. Scale bar: 100 μm.

**Figure S4.**
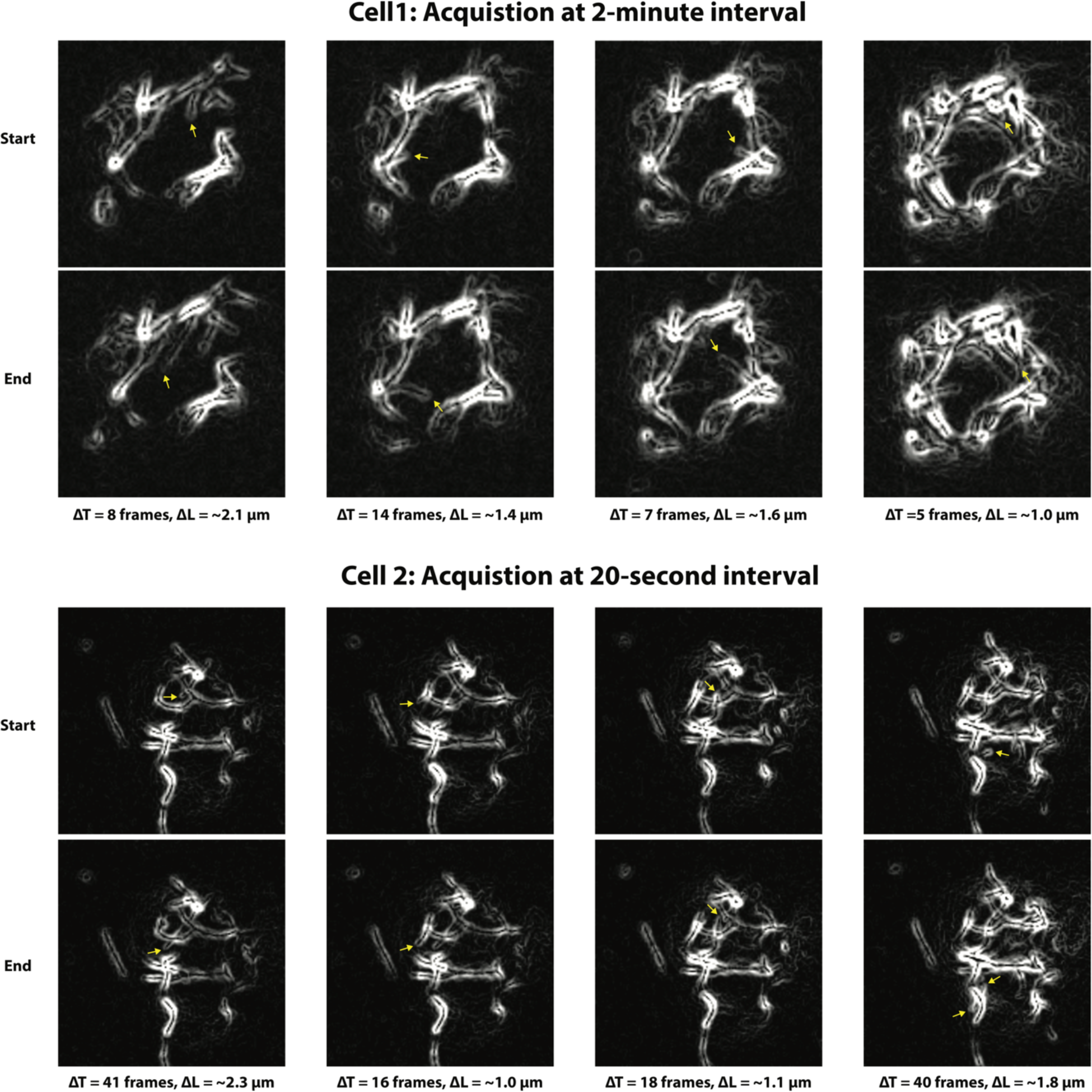
Processive elongation rate for single cellulose fibril synthesis. We estimated the TIRFM-visualized single cellulose fibrils growth rates by manually measuring the length change (ΔL) of an identifiable growing fibril between two image frames. The edge detection tool of ImageJ was used to better identify the fibril tip. Shown here are the eight image pairs from two distinct protoplast cell samples that resulted in 0.13 (± 0.04, STD) μm/min as an estimated fibril growth rate.

**Figure S5.**
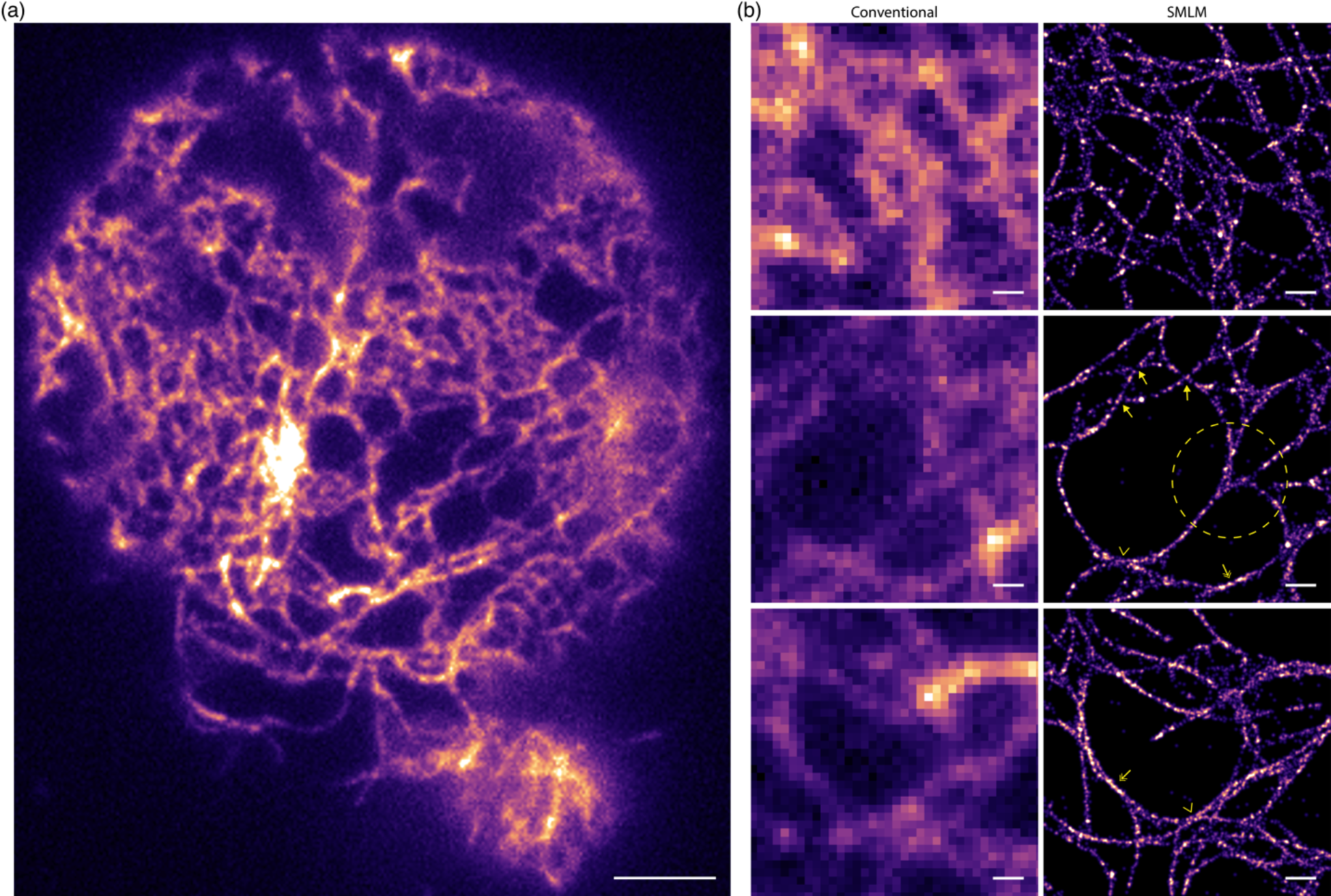
STORM imaging reveals finer ultrastructure of nascently synthesized cellulose fibrils and its assembly into a highly intertwined fibrillar network. (a) TIRFM image of a cellulose fibril network that was peeled off a protoplast cell— after cell wall regeneration for 48 hours—and adsorbed onto the poly-L-lysine-coated coverslip surface. Cellulose was labeled with Alexa568-tdCBM3a. The corresponding STORM image was presented in Figure 8A of the main text. (b) Comparative TIRFM (left panel) and SMLM/STORM (right panel) images of three selected subareas are shown here. STORM reveals the fine mesh of a reticulated network that is not apparent in TIRFM image (top). STORM enables to identify various types of inter-fibril architectures: cross-coalescence (arrow), tangential fusion of two curved fibrils (two-headed arrow), close but non-coalesced two fibrils (chevron), and multi-fibril bundling/unbundling (dotted circle). Scale bar: 5 μm (a) and 0.5 μm (b).

**Figure S6.**
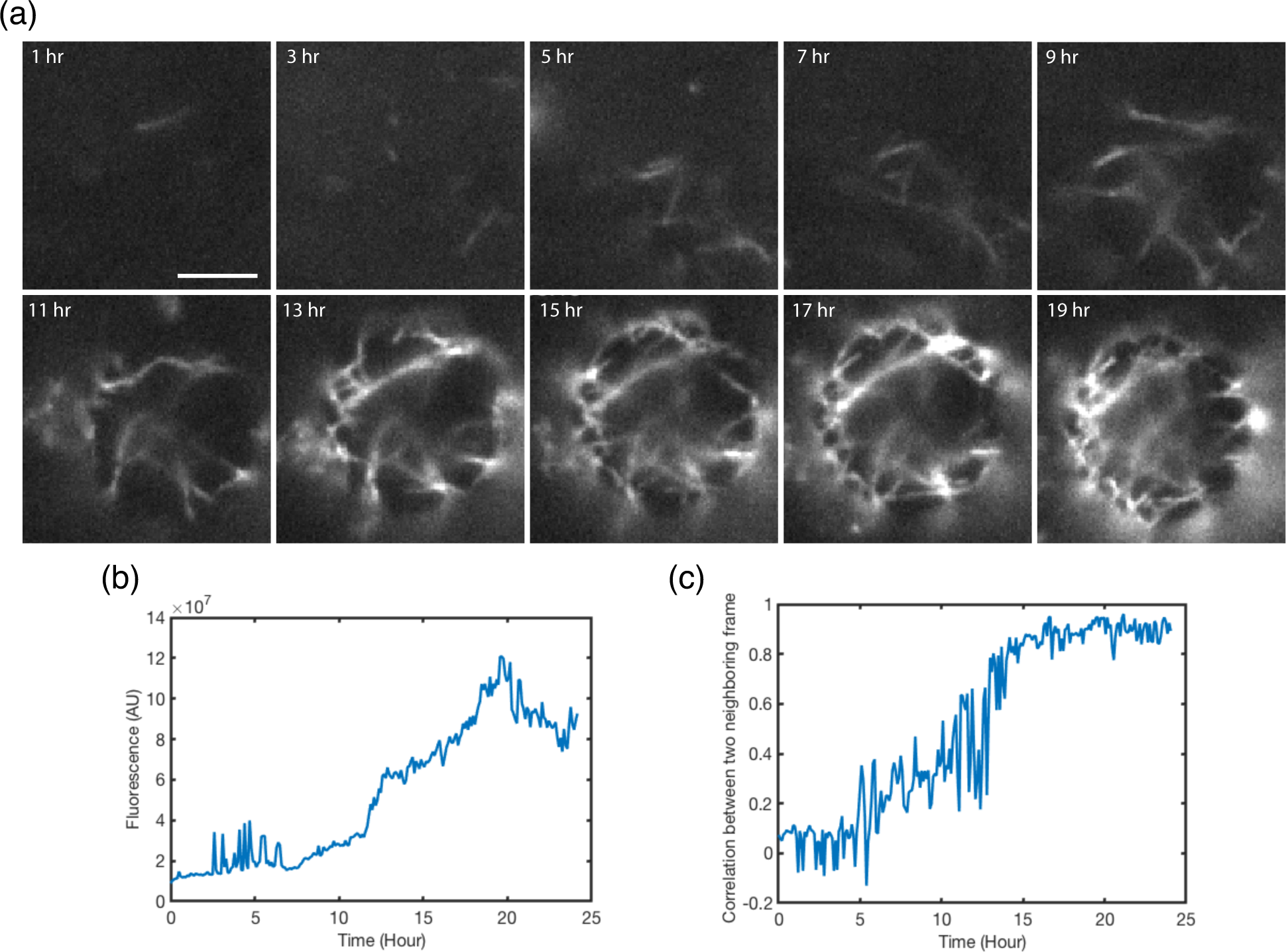
Xyloglucan knockout protoplasts show no discernible difference in cellulose fibril synthesis and assembly mechanism compared to wild-type control. (a) Time-lapse TIRFM image sequences of cellulose fibrils synthesis on a (*xxt1, xxt2*) mutant Arabidopsis protoplast surface. Cellulose was *in situ* stained with Alexa568-tdCBM3a present in the cell wall regeneration media. Image acquisition began at 2-hour after the start of cell wall regeneration and continued for ∼24 hours at 6-minute intervals. Images subsampled at 2-hour intervals are shown here (see *SI Video 11* for the full image stack). (b) Total fluorescence change in time obtained from the time-lapse fluorescence images shown in (a). (c) Normalized cross-correlation (NCC) between two consecutive image frames over time from (a-b) (see *SI Materials and Methods* for the definition) to estimate the change of overall mobility of cellulose on the protoplast surface in time. A lower NCC value implies a higher mobility and vice versa. Scale bar: 5 μm.

